# Metabolostasis failure thresholds are linked with network topology, metabolite solubility, and translational control

**DOI:** 10.64898/2026.03.31.715129

**Authors:** Shon A. Levkovich, Christine M Lim, Emi A. Marzini, Hanaa Adsi, Maoz Lahav, Ilana Sogolovsky-Bard, Myra Gartner, Keila Kaplan, Yasmin DeRowe, Metsada Pasmanik-Chor, Alexander Brandis, Michele Vendruscolo, Ehud Gazit, Dana Laor Bar-Yosef

**Author notes:** Correspondence (Michele Vendruscolo); (Ehud Gazit); (Dana Laor Bar-Yosef). Equal contribution (Shon A. Levkovich, Christine M Lim).

## Abstract

Cells maintain metabolite homeostasis (metabolostasis) by buffering fluctuations in metabolite levels, yet the limits of this buffering and the mechanisms underlying metabolic toxicity remain poorly understood. To study this, we systematically overfed metabolites in *Saccharomyces cerevisiae* and quantified associations with growth inhibition, intracellular aggregation, and multiomic perturbations. We identify metabolite-specific failure thresholds at which amyloid-like aggregates are observed, with graded growth inhibition detectable at sub-threshold concentrations, suggesting toxicity mechanisms beyond transporter saturation. Metabolites with higher network influence and broader pathway participation are associated with higher failure thresholds and smaller pathway disturbances. These patterns are associated with chemical properties and solubility: more soluble metabolites, while broadly tolerated, are associated with localised aggregates at their failure thresholds, whereas less soluble metabolites are associated with larger systemic pathway disruptions. Multiomic integration identifies a two-tiered translational regulatory architecture characterising cellular resilience to metabolic overfeeding. General resilience is associated with transcriptional commitment to resource conservation via attenuation of anabolic pathways. Metabolite-specific defense is characterised by high-magnitude translational regulatory events; for example, engagement of aromatic catabolism under phenylalanine overfeeding and energetic control pathways under glycine overfeeding. Together, our results operationally define metabolostasis as a cellular system associated with constraint of metabolite concentrations, coordination of network and pathway-level regulation, and buffering against amyloid-like aggregation, highlighting how network topology, pathway architecture, and chemical properties are associated with metabolic resilience and toxicity thresholds.

## Introduction

Cells must maintain the concentrations of hundreds of metabolites within ranges compatible with enzymatic function, network flux, and proteome integrity. This is achieved through coordinated regulation of uptake, biosynthesis, catabolism, and compartmentalization, which are processes that must respond dynamically to fluctuating nutrient availability and metabolic demand^1–3^. When this balance is disrupted, as in inborn errors of metabolism, where enzyme deficiencies lead to chronic accumulation of specific metabolites^4^, the consequences can be severe, ranging from enzymatic network dysfunction to neurological damage. Yet despite the clinical importance of metabolite dysregulation, the quantitative limits of cellular metabolite buffering capacity, and the system-level determinants of why those limits differ so dramatically across metabolites, have not been systematically characterised.

Metabolite accumulation induces toxicity via several well-established mechanisms described across bacteria, yeast, and mammalian systems. First, cross-inhibition between metabolic reactions is common as many enzymes are inhibited by structurally unrelated metabolites, thereby perturbing network-level fluxes^5,6^. Next, competition for transporters or essential cofactors can lead to systemic imbalances, for example: cysteine is a known metal ion chelator capable of altering metal homeostasis^6^. In addition, non-enzymatic reactions such as Maillard reactions between amino acids and sugars are concentration-dependent and contribute to cellular damage^7^. However, historically, these mechanisms have rarely been integrated into a unified framework of cellular metabolite management.

Emerging evidence also indicates that accumulated metabolites can self-assemble into cytotoxic, amyloid-like aggregates^8–16^, suggesting the existence of a metabolite homeostasis (metabolostasis) system that maintains metabolite concentrations within safe limits^17^. Metabolostasis encompasses coordinated enzymatic regulation, organellar storage, and compartmentalization, which together prevent metabolite aggregation and preserve cellular integrity^18,19^. For example, vacuoles in plants and fungi function as repositories for potentially toxic metabolites^20^, and mitochondrial-derived compartments facilitate adaptation to excess amino acids by promoting their catabolism^21^.

While metabolostasis mechanisms can buffer minor fluctuations in metabolite levels, acute disturbances may overwhelm the system, leading to toxicity. The toxic effects of amino acid excess were first documented in bacteria and yeast in the mid-20th century^22–24^, and studies in humans suggest upper limits for amino acid intake, with excessive consumption linked to neurological symptoms^25–27^. However, cellular responses to acute metabolite overload, particularly system-level constraints that determine tolerance and failure, remain largely unexplored.

Adaptation to metabolite overload involves complex cell-wide programs. These adaptive responses involve regulatory programs that act at multiple levels including but not limited to transcription. Integrative analyses have demonstrated that while most often used as a proxy for functional activity, transcript levels explain only a limited fraction of protein-level variance^28,29^, underscoring the importance of integrating regulatory effects beyond transcription. For example, translational efficiency, protein turnover, and other regulatory mechanisms contribute to proteome composition independently of transcript abundance^30^. It has been reported that cells post-transcriptional buffering occurs to stabilize protein levels despite transcript fluctuations especially under stress conditions^31^. Given this, it is necessary to employ a system-wide approach to better characterise adaptive responses modulating cellular tolerance and stress resilience.

Here we systematically characterise metabolostasis through controlled metabolite overfeeding in *Saccharomyces cerevisiae*, quantifying failure thresholds for twenty-three metabolites alongside metabolomic, transcriptomic, and proteomic responses. While single-metabolite perturbations do not fully recapitulate the complexity of physiological metabolite mixtures, this framework enables systematic mapping of metabolite-specific buffering capacity and exposes failure modes that may be obscured under combinatorial conditions. Integrating these data, we find that failure thresholds are associated with metabolic network topology and metabolite solubility, and that cellular resilience is characterised by a two-tiered post-transcriptional regulatory architecture, which emerges as a general resource-conservation program superimposed by metabolite-specific high-magnitude translational responses. Amyloid-like aggregates are observed coincidentally with growth inhibition at failure thresholds, consistent with aggregation representing a feature of metabolostasis failure, although its causal contribution to toxicity remains to be established. Together, these findings provide a quantitative framework for understanding metabolic resilience and its limits, with potential relevance to conditions characterised by chronic metabolite accumulation including inborn errors of metabolism.

## Results

### Metabolite overfeeding defines failure thresholds for cellular tolerance

To investigate the limits of metabolite buffering, we systematically overfed 20 amino acids and 4 nucleobases to *Saccharomyces cerevisiae*. Guanine was excluded due to low solubility. Dose-response growth measurements **(Figure 1A)** revealed metabolite-specific failure thresholds **(Figure 1B)**, above which overfeeding caused sharp reductions in growth, often starting below transporter saturation, indicating limits in cellular regulatory buffering rather than uptake **(Figure 1C)**. Failure thresholds varied widely **(Figure 1B)**: cysteine was the most toxic (∼1.3 mM), followed by phenylalanine, threonine, and isoleucine (∼11–12 mM), while lysine, glutamic acid, and proline were tolerated above 1 M. Several metabolites, including tyrosine, aspartic acid, uracil, cytosine, leucine, histidine and adenine, did not have significant dose-dependent toxicity and did not reach definable failure thresholds under the conditions tested **(Figure S1)**. Tyrosine is extremely insoluble in water, and thus the maximal concentration that could be examined was 2 mM. To exclude non-metabolite specific effects on the observed toxicity, we examined the conditions at which osmotic pressure (sorbitol treatment), excess carbon source (glucose treatment), and excess nitrogen (ammonium sulfate treatment) affect growth **(Figure S2)**. Our control experiments confirmed that failure thresholds below 400 mM reflected metabolite-specific effects rather than general stress induced toxicity **(Figure S2)**. Here, we establish quantitative failure thresholds for metabolite tolerance and provide a foundation for exploring mechanistic consequences, including aggregation and pathway perturbations. Thus, for the rest of this work, we focused on nine amino acids as our core metabolite set: eight amino acids with the lowest failure thresholds (cysteine, phenylalanine, threonine, isoleucine, alanine, methionine, tryptophan, and valine), as well as, glycine the simplest amino acid that does not possess either aliphatic or aromatic side chain. Recently, a phenylalanine (Phe) salvage model was established, showing that toxicity correlates with Phe accumulation and aggregation can be observed already at 0.9 mM (following mutation in Aro4 enzyme that takes part in the shikimate pathway), whereas WT cells can tolerate this concentration^32^. Accordingly, the amount of Phe required to induce toxicity in the salvage model is significantly lower than in WT cells. These results imply that toxicity arises from elevated Phe levels and self-assembly, either due to high external supplementation or genetic manipulation leading to intracellular accumulation, suggesting that the failure threshold is unlikely to be explained solely by transporter saturation **(Fig. 1C)**.

**Figure 1.**
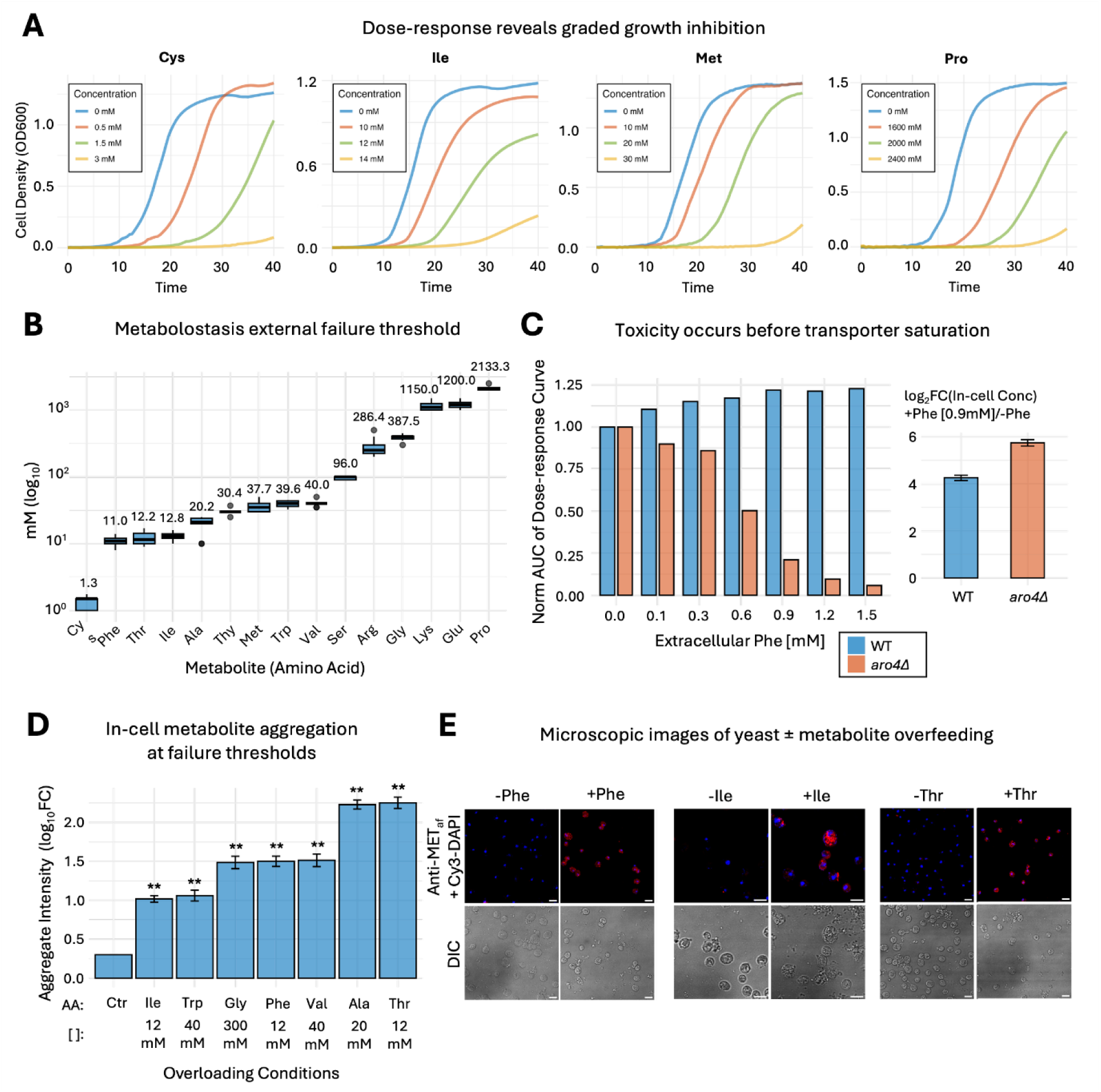
Dose-dependent metabolite overfeeding reveals failure thresholds and coincident aggregation in yeast. **(A)** Representative dose–response growth curves illustrating metabolite-specific tolerance limits. **(B)** Metabolite-specific failure thresholds. Failure thresholds were estimated from dose-response growth measurements using the linear-regression procedure described in Methods, revealing broad variation across metabolites, from highly toxic cysteine (∼1.3 mM) to well-tolerated lysine, glutamic acid, and proline (>1 M). Several metabolites did not reach definable thresholds under the conditions tested. **(C)** Growth inhibition is dose-dependent as reflected by decreasing area-under-the-curve (AUC) of yeast growth curve with increasing Phe feeding in the *aro4*Δ yeast strain. Because growth defects emerge at concentrations below the estimated failure threshold, the limitation is unlikely to be explained solely by transporter saturation. In cell concentration is calculated as +Phe/-Phe ratio in the presence of 0.9 Mm Phe. The data presented here were derived from the analysis of raw datasets reported in previous studies^32^. **(D)** Amyloid-like aggregate formation at metabolite-specific failure thresholds using the amyloid-specific dye ProteoStat. Error bars represent standard deviation (SD) from the mean. **(E)** Representative confocal images illustrating intracellular aggregate signals at concentrations near the failure thresholds of the designated metabolites, following immunostaining with antibodies raised against fibrils of the same metabolites, designated as Anti-METaf (=amyloid fibrils). Overlay with DAPI is shown for all fluorescent images (Cy3-DAPI). The scale bar is 5 µm.

### Metabolite aggregation coincides with metabolite-specific failure

Overfeeding yeast with metabolite concentrations at the metabolite-specific failure thresholds triggers the formation of intracellular amyloid-like aggregates. To study the association of aggregation with metabolite toxicity, we quantified intracellular amyloid-like assemblies upon metabolite overfeeding **(Figure 1D)** using the amyloid-specific dye ProteoStat, previously been reported to stain metabolite aggregates in vivo^9^. A significant increase in aggregate signal was observed at the failure threshold concentrations of several amino acids, with up to a ∼100-fold increase in fluorescence depending on the metabolite **(Figure 1D)**. Since ProteoStat could also potentially stain co-aggregating proteins, we also performed immunostaining with preparations of antibodies raised specifically against each type of metabolite assemblies for the three structurally diverse amino acids that have been shown to self-assemble: phenylalanine^8^, isoleucine^14^, and threonine **(Figure S3)**. Staining with the metabolite-specific antibodies (either against phenylalanine-, isoleucine-, or threonine-assemblies) detected distinct intracellular extranuclear puncta at the treated failure thresholds **(Figure 1E)**. These results are consistent with specific aggregation processes occurring in association with metabolostasis failure at threshold concentrations of each metabolite in turn.

### Metabolic signatures of overfeeding

To quantify how individual metabolites reshape the metabolic landscape, we profiled intracellular amino acid concentrations at sub-toxic supplementation levels (∼50-60% of each metabolite’s failure threshold). Across all conditions, cells displayed a marked increase in the intracellular concentration of the overfed metabolite **(Figure S4)**, confirming robust uptake even under impending toxicity. Beyond these primary effects, each amino acid induced a distinct metabolic signature **(Figure S4)**, as revealed by targeted LC–MS/MS profiling. While the global structure of the metabolome remained highly reproducible across replicates, the direction and magnitude of metabolite-induced shifts were strongly condition-specific, indicating coordinated rewiring of metabolic regulation **(Figure S5A)**. Notably, many perturbations triggered consistent depletion of arginine and ornithine **(Figure S5B)**, suggesting that the urea cycle may be implicated as a recurrent buffer of acute metabolic imbalance^33^.

### Network topology and pathway engagement determine metabolite tolerance

To understand system-level features (metabolomic level) determining metabolite failure thresholds, we next quantified how each metabolite perturbs other metabolites upon overfeeding. Using causal metabolic perturbation data, we (1) set up the metabolic regulatory network **(Figure 2A)** and (2) computed a metabolite-specific regulatory influence score that captures how strongly each metabolite affects the levels of other metabolites **(Figure 2B)**. Details on defining the metabolic regulatory network and calculating the regulatory influence score are described in **Methods**. Strikingly, we find that failure thresholds were positively correlated with network influence **(Figure 2C)**, indicating that metabolites exerting broader regulatory control are also those the cell can tolerate at higher concentrations. Classification of metabolites based on their global regulatory influence (‘Central’ as PPR > mean (PPR) and ‘Peripheral’ as PPR < mean (PPR)) reveals that central metabolites have higher failure thresholds **(Figure 2D)**. Metabolites with greater pathway participation showed similarly elevated failure thresholds **(Figure 2E)**, suggesting that integration into multiple pathways provides a distributed buffering architecture. Together, these two findings **(Figure 2D, E)** suggest that metabolites with higher regulatory influence and pathway involvement are likely to be better buffered, thus the system can withstand higher perturbations before failure is reached (higher failure thresholds). Consistent with this, a heatmap of relative pathway disturbances **(Figure 2F)** revealed that low-threshold metabolites caused disproportionately large pathway perturbations, whereas high-threshold metabolites induced smaller, more localized changes **(Figure 2G)**. These results demonstrate that system-level features of the metabolic network - regulatory influence and pathway connectivity - govern the capacity of the metabolostasis system to absorb acute metabolite excess and determine the onset of toxicity.

**Figure 2.**
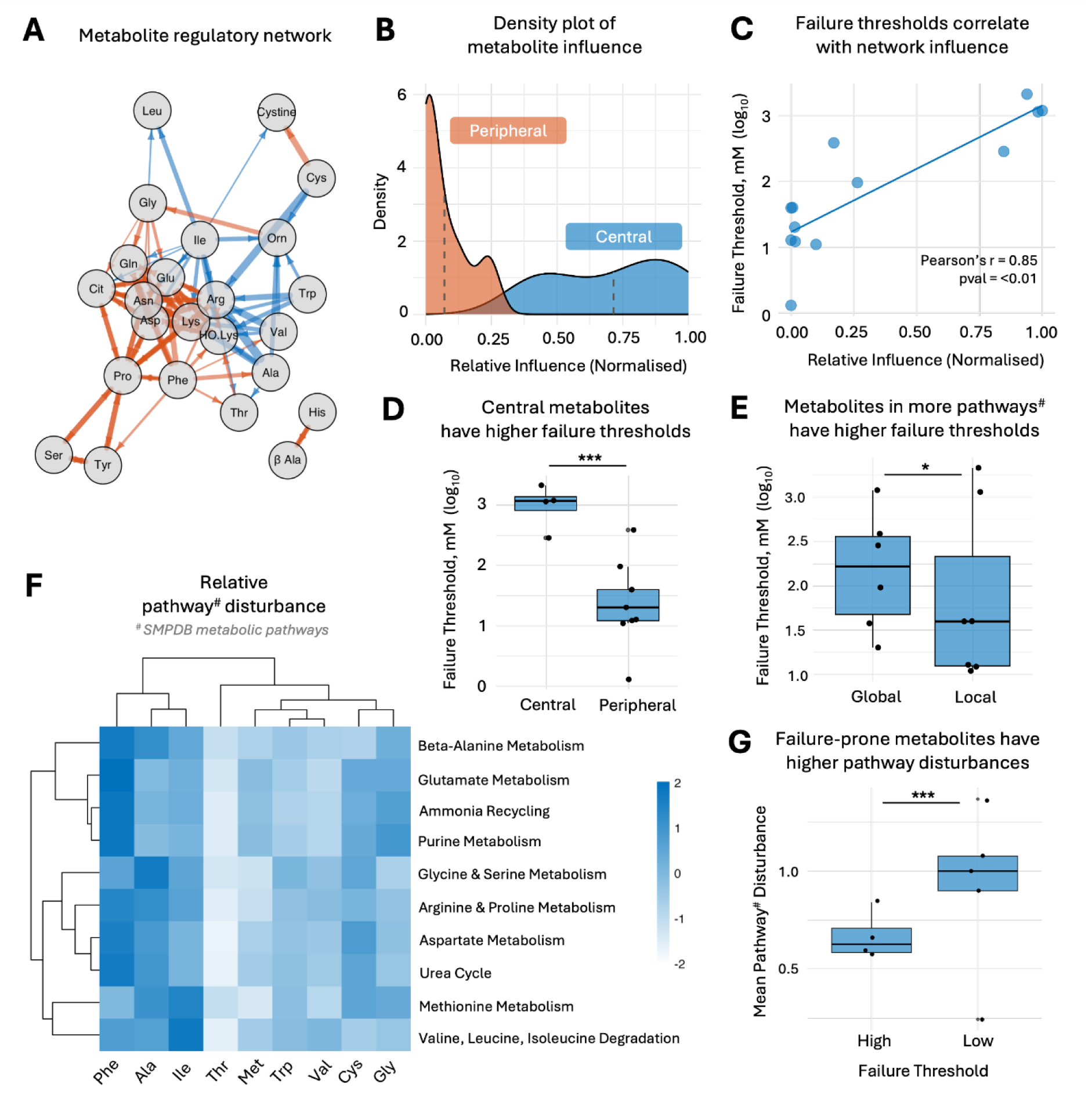
Metabolic network topology and pathway connectivity are associated with metabolite-specific failure thresholds and systemic perturbations. **(A)** Metabolic regulatory network constructed from: (1) directed perturbation edges from overfed metabolites to significantly altered metabolites, and (2) strong correlations among the remaining metabolite pairs. **(B)** Density distribution of metabolite regulatory influence, quantified by the personalized PageRank (PPR) score of each metabolite within the regulatory network. **(C, D)** Network regulatory influence is positively associated (r = 0.85, p < 0.01) with failure thresholds (C), and metabolites with higher regulatory influence (central) exhibit correspondingly higher failure thresholds (D), consistent with enhanced buffering capacity. **(E)** Metabolites participating in a greater number of pathways are associated with higher failure thresholds prior to growth inhibition. **(F)** Heatmap depicting relative pathway disturbances following metabolite overfeeding. Rows represent individual metabolites, and columns represent metabolic pathways. **(G)** Metabolites with higher failure thresholds are associated with smaller pathway perturbations, whereas low-threshold metabolites are linked with broader pathway disruptions. Statistical significance was calculated using a Wilcoxon test; (*) p ≤ 0.05 and (***) p ≤ 0.001.

### Metabolite chemical properties & solubility govern systemic impact and aggregation

We next explain how chemical properties and solubility are fundamental predictors of metabolite perturbation effects and aggregation susceptibility. We first establish that chemical properties determine solubility **(Figure 3A)**. When analyzing the tested metabolites, we find that chemical groups with higher solubility partake in more pathways (are more global) **(Figure 3B)** and cause lower pathway disturbance **(Figure 3C)** upon overfeeding. From these trends, we infer that metabolites with higher solubility are more functionally relevant, and are hence more buffered by the cellular system, producing weaker global network disturbances. In contrast, low-solubility metabolites triggered larger metabolic perturbations, indicating that poor solubility amplifies system-wide stress. To support this hypothesis, we find that solubility correlates strongly with baseline intracellular abundance **(Figure 3D)**, revealing that cells are naturally able to maintain higher steady-state levels of more soluble metabolites and may have evolved regulatory architectures tuned to their physicochemical properties. In addition, metabolite solubility is negatively correlated with aggregation propensity – more soluble metabolites tend to have lower aggregation propensities **(Figure 3E)**. These findings demonstrate that chemical properties and solubility dictate how cells distribute and maintain metabolite levels, thereby constraining cellular tolerance to metabolic perturbation and aggregation susceptibility^34^. To provide a mechanistic explanation for the failure thresholds across metabolites, we investigated the correlation between the experimentally observed failure thresholds and various biophysical properties of the metabolites. Notably, we observed a significant correlation in the case of hydrophobicity and β-sheet propensity of metabolites (**Figure 3F**), suggesting that the toxicity of metabolites is associated with their aggregation propensity. This is confirmed by the strong correlation between aggregation propensities and failure thresholds even at different pH **(Figure 3G)**.

**Figure 3.**
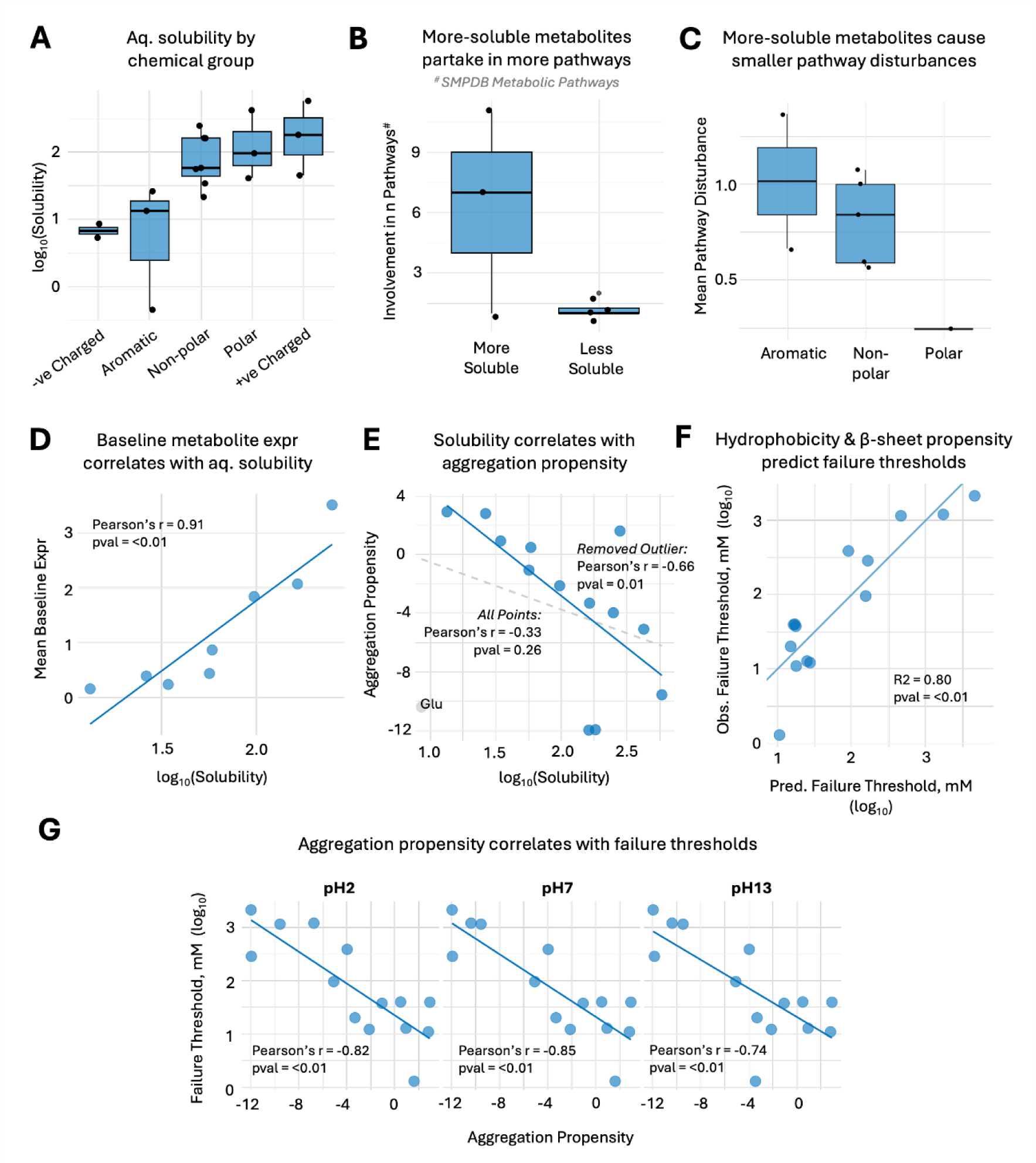
Metabolite solubility predicts pathway integration, intracellular abundance, and aggregation levels. **(A)** Chemical properties govern metabolite solubility. **(B,C)** More soluble metabolites partake in more pathways (B) and cause lower pathway disturbance (C). These functional consequences show that soluble metabolites are likely to be more functionally relevant and are hence better buffered by the metabolostasis system to ensure resilience. **(D)** A positive correlation (r = 0.91, p < 0.01) between baseline intracellular abundance and solubility suggests that cellular metabolite pools are tuned to the physicochemical properties of individual metabolites. **(E)** Metabolite solubility is negatively correlated with aggregation propensity (r=-0.66, p = 0.01). **(F)** Polynomial regression model predicts experimentally observed failure thresholds from metabolite hydrophobicity and β-sheet propensity. The model showed a significant overall fit (F = 8.99, p < 0.01, R² = 0.80). Hydrophobicity exhibited a significant quadratic relationship with failure threshold (p < 0.05), whereas β-sheet propensity was not significant. Hydrophobicity and β-sheet propensity of each metabolite is available in **Table S2**. **(G)** Metabolite aggregation propensity negatively correlates with failure thresholds across pH2, 7 and 13, where metabolites that are more prone to aggregation have lower failure thresholds.

### T:P Control Reveals Two Distinct Regimes for Metabolic Resilience

To investigate regulatory principles governing metabolic tolerance, we study the post-transcriptional (T:P) regulatory landscape. For this, we introduce the Adaptive Strategy Preference (ASP) score **(Methods)** that quantitatively represents the relative preference for adaptive attenuation (resource conservation) versus adaptive gain (rapid protein synthesis). The protein and RNA were obtained from the same culture, thus enabling the reliable comparison of the omic results. Across all metabolite overfeeding conditions, the ASP score exceeded 1 **(Figure 4A)**, indicating that adaptive attenuation - a strategy of repressing non-essential genes to conserve resources - remains the predominant adaptive strategy under metabolic stress.

**Figure 4.**
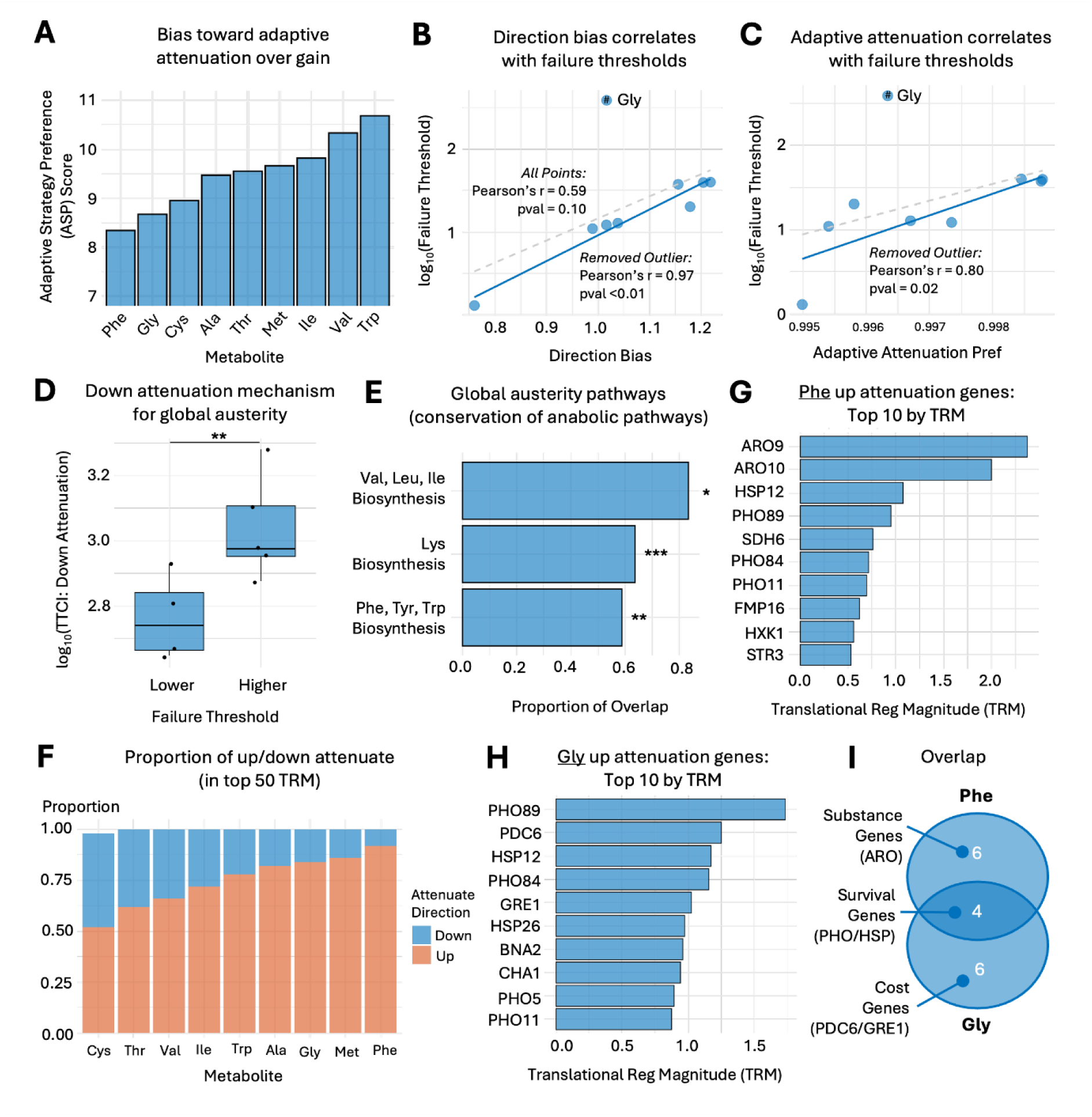
Translational control identifies a two-tiered regulatory architecture associated with metabolic resilience. **(A)** Adaptive Strategy Preference (ASP) score quantifying the relative preference for adaptive attenuation over adaptive gain across metabolite overfeeding conditions. **(B)** Directional bias correlates with core resilience. The T:P directional bias correlates (r = 0.97, p < 0.01) with failure thresholds across the core metabolite set (the nine lowest-threshold metabolites), consistent with resource conservation as a dominant global resilience strategy. **(C)** Adaptive Attenuation Preference is positively associated with resilience (r = 0.80, p = 0.02), consistent with a greater reliance on attenuation-based buffering under metabolite stress. **(D)** Metabolites with higher failure thresholds are associated with significantly greater down-attenuation TTCI (Wilcoxon test, p < 0.05). **(E)** Down-attenuated genes conserved across more than half of the nine overfeeding conditions are significantly enriched for costly anabolic pathways, including amino acid biosynthesis, consistent with a global resource-conservation strategy. **(F)** Up-attenuation predominates among high-intensity regulatory responses. Ranking genes by translational regulatory magnitude (TRM; top 50) shows that the largest-amplitude regulatory events are predominantly up-attenuation, consistent with high-intensity, metabolite-specific defense programs (as reflected by TTCI expenditure). **(G)** (Under Phe overfeeding, prominent targets include aromatic catabolism genes (*ARO9/ARO10*), consistent with induction of rerouting pathways that may limit toxic intermediate accumulation. **(H)** Under glycine overfeeding, prominent targets include genes linked to energetic control (e.g., PDC6) and cytoprotection (e.g., GRE1, HSP26). **(I)** Overlapping up-attenuated genes across overfeeding conditions are enriched in core stress-response and survival pathways, consistent with a shared resilience infrastructure.

Analysis of the core metabolite set revealed a multi-layered defense. We find an extremely strong positive correlation (r = 0.97, pval < 0.01) between directional bias – measuring the overall orientation of post-transcriptional regulation toward repression versus induction **(Methods)** – and failure thresholds **(Figure 4B)**. This suggests that effective repression is the primary determinant of resilience over total induction as a system-wide strategy of resource conservation to maximize the repression of non-essential genes. Similarly, the Adaptive Attenuation Preference (AAP) score – proportion of genes exhibiting attenuation relative to all significantly perturbed genes **(Methods)** – also showed a significant positive correlation (r = 0.80, pval < 0.05) **(Figure 4C)**. This reflects that general resilience of the cell to metabolite toxicity is successfully predicted by both the global transcriptional commitment to conservation and the frequency of successful T:P buffering events.

We note that Glycine’s ultra-resilience acts as an outlier and is not well explained by the occurrence of successful buffering events. Hence, we hypothesize that the intensity of regulatory response may enable enhanced tolerance to metabolic toxicity. To test this, we introduced the Total T:P Control Intensity (TTCI) metric **(Methods)**, quantitatively representing the total regulatory budget the cell spends to achieve survival under stress. We find that metabolites with higher failure thresholds (higher resilience) exhibits a significantly greater down-attenuation TTCI **(Figure 4D)**, demonstrating that enhanced-resilience requires a larger commitment to regulatory intensity (preparedness).

Based on our data, we propose a two-tiered regulatory mechanism governing metabolostasis resilience upon metabolite overfeeding: general resilience toward toxicity, and metabolite-specific buffering. General resilience is governed by global austerity, which is the conservation of resources. At a functional level, over-representation analysis of consistently down-attenuated genes across all treatments (down-attenuated in more than half of all metabolite overfeeding conditions) showed significant enrichment for costly anabolic pathways (amino acid biosynthesis) **(Figure 4E)**. Repression of these pathways ensures the generation of the necessary pool of free resources required for systemic defense. Metabolite-specific stressors are then more precisely buffered by high-magnitude regulatory events (high TRM up-attenuation) **(Figure 4F)**. We exemplify the functional targets of this metabolite-specific defense for Phe (low failure threshold, low resilience) **(Figure 4G)** and Gly (high failure threshold, high resilience) **(Figure 4H)**. The presence of overlapping genes demonstrates the activation of common stress response pathways (PHO genes and HSPs) **(Figure 4I)**, while metabolite-specific targets reveal that Phe-specific response occurs via promoting aromatic catabolism (Aro9/Aro10) to counter the rise in toxic flux, and promoting energetic control and cytoprotection in response to Gly overfeeding. To test this hypothesis, we compared the specificity of Aro(s), Pho(s) and Hsp(s) and expected more metabolite-specific buffering (Aro(s)) to be involved in buffering across fewer metabolite overfeeding conditions than common stress response pathways (Pho(s) and Hsp(s)). Our results confirm this, where metabolite-specific buffering genes are involved in buffering fewer metabolites under overfeeding conditions: Aro(s) buffer on average 2.22; Pho(s) 4.8; and Hsp(s) 3.8 metabolites (of 9) under overfeeding conditions **(Methods)**.

Moving forward, we test our hierarchical buffering model, by first examining metabolite-specific adaptive rerouting mechanisms, followed by interrogating shared stress infrastructure pathways.

### Adaptive metabolic rerouting (ARO genes) buffers toxic flux

Findings from our T:P analysis identify ARO genes contribute to buffering of toxicity effect of Phe overfeeding. To determine whether these modules are functionally required for resilience, we examined growth under Phe overfeeding in WT, aro10Δ, and aro4Δ strains, representing distinct components of the aromatic amino acid metabolic network (**Figure 5A**). Exposure to 8 mM Phe induced significant growth inhibition in WT cells. This toxicity was markedly exacerbated in both aro10Δ and aro4Δ strains (**Figure 5B**), demonstrating that intact aromatic amino acid metabolic pathway function is required for optimal tolerance to Phe stress (**Figure S6**). Importantly, Aro10-dependent buffering exhibited a threshold-like phenotype. Loss of Aro10 did not significantly alter growth at basal or moderate Phe concentrations but resulted in pronounced sensitivity at high Phe levels (**Figure 5C, D**). This concentration-dependent requirement supports a model in which Aro10 functions as an adaptive buffering mechanism engaged only once metabolite burden surpasses baseline homeostatic capacity. In contrast, aro4Δ cells displayed impaired growth even at moderate Phe concentrations (**Figure 5C, D**), consistent with disruption of upstream shikimate pathway regulation and intracellular Phe accumulation and aggregation^32^. Thus, while Aro4 contributes to general metabolic homeostasis, Aro10 mediates stress-contingent adaptive detoxification. Together, these results distinguish constitutive pathway integrity from inducible adaptive buffering within the aromatic amino acid network.

**Figure 5.**
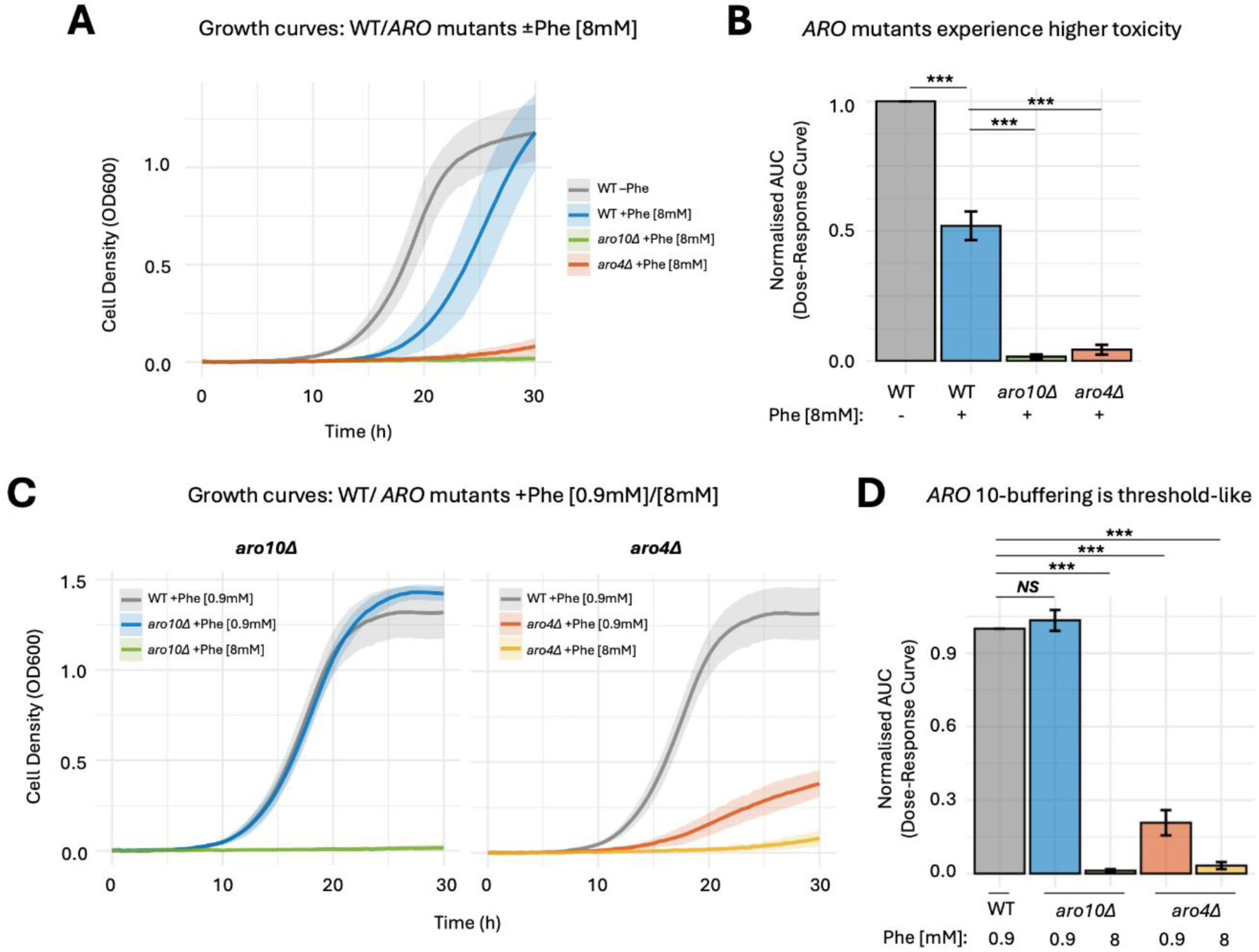
The two-tiered regulatory pattern consists of a specific adaptive buffering module and a general resilience infrastructure, as explained by the aromatic amino acid pathway (A-D) and the PHO pathway (E-F). **(A)** Growth curves of WT, *aro4*Δ, and *aro10*Δ yeast strains treated with Phe [8 mM]. Phe supplementation reduces growth in WT cells, an effect further exacerbated in *aro10*Δ and *aro4*Δ strains, consistent with a requirement for the aromatic amino acid pathway function for optimal tolerance to Phe stress. **(B)** Toxicity was quantified by calculating the area under the curve (AUC) for each strain ± Phe and normalizing values to WT under untreated conditions (−Phe). **(C)** Growth curves of *aro4*Δ and *aro10*Δ strains under moderate (0.9 mM) versus high (8 mM) Phe supplementation. *aro10*Δ cells show growth comparable to WT at 0.9 mM Phe but display pronounced sensitivity at 8 mM. In contrast, *aro4*Δ cells exhibit growth impairment that is evident at moderate Phe and increases further at high Phe. **(D)** Loss of Aro10 is associated with a threshold-dependent growth defect, consistent with an adaptive buffering role that becomes limiting only under high Phe burden. Loss of the upstream modulator Aro4 is associated with broader sensitivity across Phe concentrations. Statistical comparisons were performed using ANOVA followed by Dunnett’s test; (***) p ≤ 0.001 and *NS*, not significant. Error bars represent standard deviation (SD) from the mean.

In addition to metabolite-specific adaptive rerouting (Aro10-mediated), our model also reasons that shared preparedness pathways provide a foundational resilience layer across metabolite stresses **(Figure 4I)**.

### Polyphosphate provides a physicochemical defense by suppressing metabolite self-assembly

Polyphosphate (polyP) is a universally conserved biopolymer that influences a wide spectrum of cellular processes, including phosphate storage, osmotic and pH homeostasis and stress adaptation^35^. Since *PHO* genes were recurrently activated across metabolite overfeeding conditions **(Figure 4I)**, we hypothesized that polyphosphate metabolism constitutes the general resilience infrastructure. To test this, we examined the effect of disrupting PHO components on tolerance to metabolite overload, using Phe as a representative stressor. We compared growth of WT, *vtc4*Δ (polyP polymerase), *pho81*Δ (PHO pathway regulator)^35,36^, and *pho84*Δ (high-affinity phosphate transporter)^37,38^ under Phe overfeeding. Exposure to Phe [8 mM] significantly reduced growth in all strains. However, polyP-deficient mutants exhibited markedly greater sensitivity relative to WT **(Figure 6A & Figure S7)**. These findings demonstrate that polyphosphate metabolism is required for effective tolerance to Phe overload.

**Figure 6.**
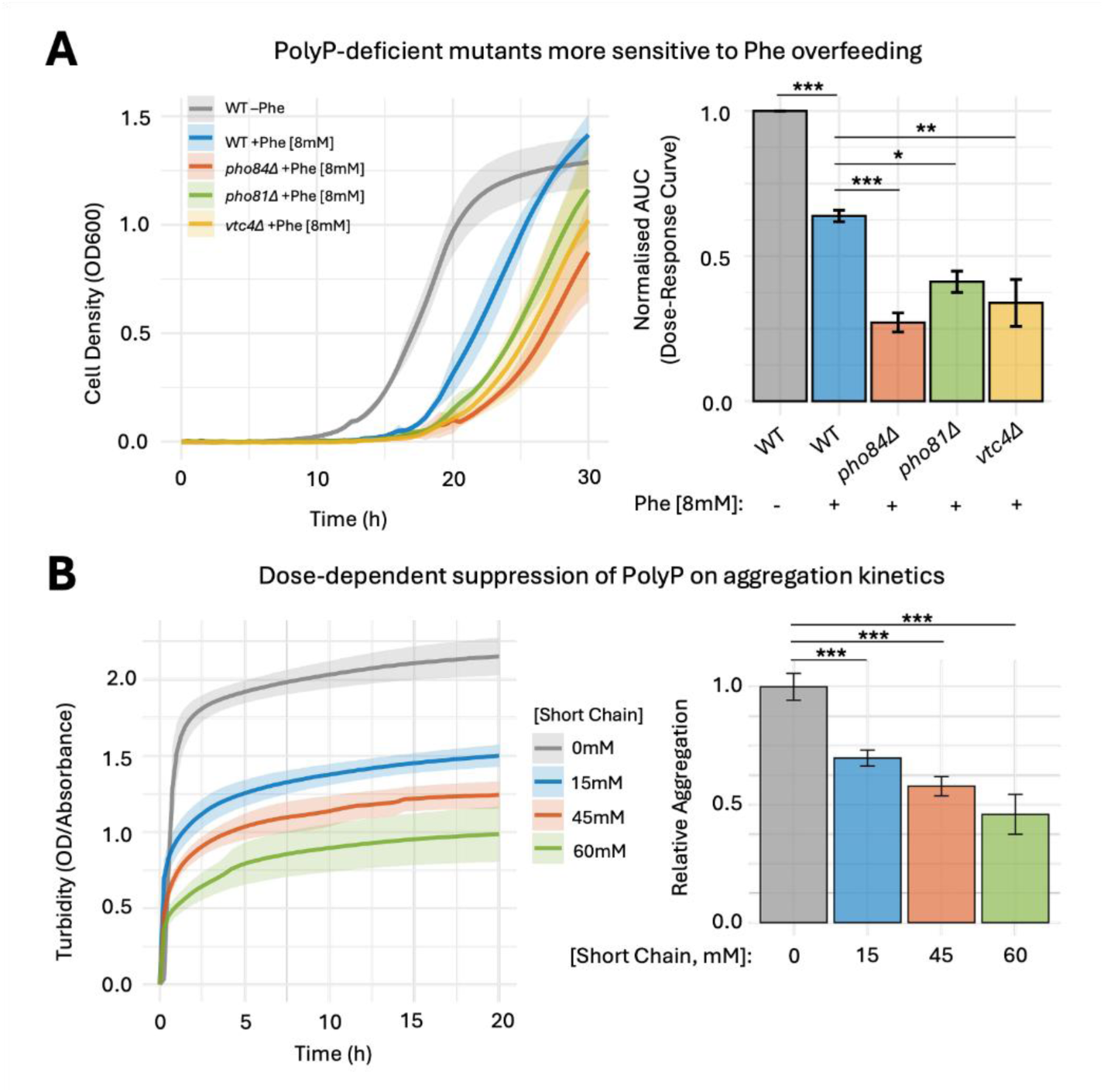
PHO pathway–mediated polyphosphate metabolism functions as a general resilience infrastructure within the hierarchical metabolostasis model. **(A)** Growth curves of WT, *pho84*Δ (high-affinity phosphate transporter), *pho81*Δ (PHO pathway regulator), and *vtc4*Δ (polyphosphate polymerase) strains under Phe overfeeding (8 mM). Phe supplementation reduces growth across all genotypes, with polyphosphate-deficient mutants exhibiting significantly greater sensitivity than WT. This is consistent with a role for PHO pathway-dependent polyP metabolism in tolerance to Phe overload, supporting the general preparedness layer of the metabolostasis model. **(B)** In vitro aggregation kinetics of Phe in the presence of increasing polyphosphate (polyP) concentrations. Phe alone shows a time-dependent increase in turbidity, whereas addition of polyP suppresses turbidity in a dose-dependent manner, with 60 mM polyP producing the strongest inhibition. Endpoint turbidity quantification indicates a 30-50% reduction in endpoint turbidity across increasing polyP concentrations. Statistical comparisons were performed using ANOVA followed by Dunnett’s test; (*) p ≤ 0.05, (**) p ≤ 0.01, (***) p ≤ 0.001.

PolyP has been demonstrated to interact with mature amyloid fibrils enriched in β-sheet structures and decrease the generation of toxic oligomeric intermediates^35,39^. Thus, we hypothesize that polyP may influence the physicochemical landscape of metabolite toxicity^40^. To study this, we measured aggregation kinetics in vitro in the presence of increasing polyP concentrations **(Figure 6B)**. Phe alone displayed rapid and sustained turbidity increases. Addition of polyP resulted in a dose-dependent suppression of aggregation kinetics, with polyP [60 mM] being most inhibitory. Quantification of endpoint turbidity revealed ∼30–50% reduction in aggregation across increasing polyP concentrations. These results demonstrate that disruption to phosphate metabolism compromises effective tolerance to metabolite stress.

Collectively, our findings indicate that polyphosphate metabolism contributes to intracellular resilience by sustaining metabolic resource buffering and attenuating metabolite self-assembly, functionally supporting the general preparedness layer of the metabolostasis model.

### Combinatory effects of metabolite toxicity

We further explore the interaction effects of metabolites when treated in combination. For this, we assessed yeast growth in the presence of single vs. pairwise treatments **(Figure 7A)**. Combinatory effect was studied for three structurally diverse amino acids (phenylalanine, isoleucine, and threonine) that were shown in this study to have intracellular metabolite assemblies both using the amyloid-dye as well as specific antibodies **(Figure 1E)**. In total, three combinations were tested: Phe [4 mM] + Ile [4 mM], Phe [4 mM] + Thr [4 mM], and Thr [4 mM] + Ile [4 mM]. Bliss independence^41^ was used to compute the expected fractional inhibition of a combination of two metabolites based on the individual effects of each metabolite’s overfeeding alone. Our results reveal that treatment with all metabolite pairs tested resulted in a larger growth inhibition compared to expected inhibition rate assuming simple additive effect between single metabolite treatments **(Figure 7B)**. This result indicates that metabolite toxicity is synergistic for Phe, Ile, and Thr.

**Figure 7.**
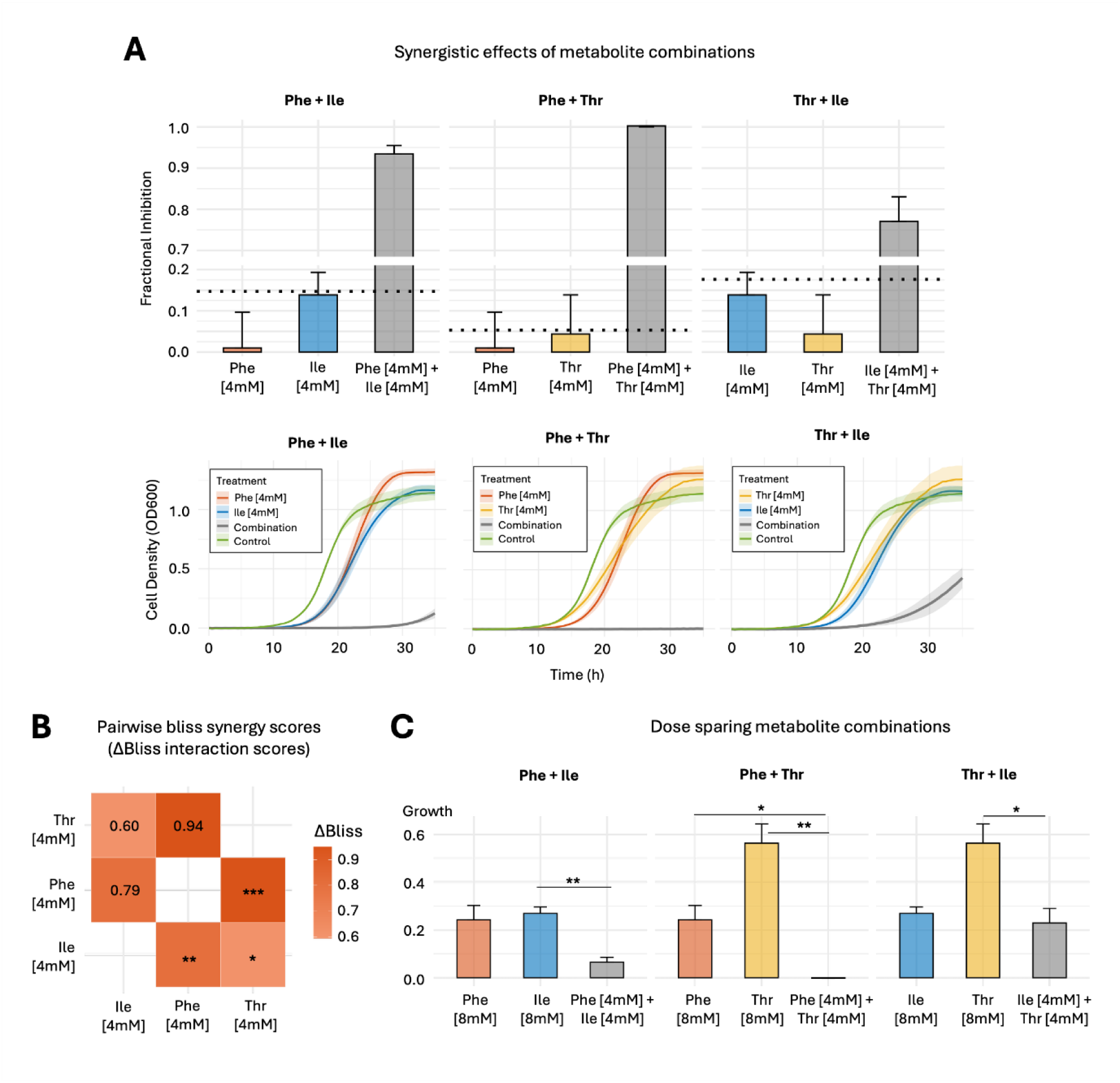
Pairwise metabolite combinations are associated with synergistic growth inhibition. **(A)** Fractional inhibition of pairwise amino acid combinations (Phe + Ile, Phe + Thr, and Thr + Ile; 4 mM each) exceeds the inhibition predicted under Bliss independence (horizontal dotted line), consistent with synergistic interactions between metabolites at sub-threshold concentrations. Fractional inhibition was calculated as AUC_treatment_/AUC_control_. **(B)** Pairwise Bliss synergistic scores for metabolite combinations of [4 mM]. The one-sample t-test with Benjamini-Hochberg multiple testing correction was used to calculate significance: (***) pval < 0.01 and (**) pval < 0.05. **(C)** At the same total metabolite concentration (8 mM), pairwise combinations (4 mM + 4 mM, 1:1 ratio) are associated with greater growth inhibition than either constituent metabolite administered alone at 8 mM. Bars represent mean ± s.e.m. Statistical significance was tested using the Welch two-sample t-test with BH multiple testing correction; (*) p ≤ 0.05 and (**) p ≤ 0.01.

To study dose sparing, we compare metabolite combinations (1:1 ratio) to each of the constituent metabolites at the same overall combination. We find that pairwise treatment of Phe with either Ile or Thr to a total of 8 mM ([4 mM] each) resulted in a larger extent of growth inhibition compared to single metabolite treatment of 8 mM **(Figure 7C)**. Notably, the combination of Phe and Thr produced a synergistic phenotype that ultimately resulted in lethality. Moreover, in agreement with the data presented in **Figure 1C**, these results support that the observed toxicity does not arise from simple transporter saturation but rather from a synergistic effect whose emergent properties exacerbate cellular stress and suppress yeast growth.

## Discussion

Maintaining cellular homeostasis requires that metabolite concentrations remain within ranges compatible with growth and function. Here, we characterised metabolostasis as a multi-layered system whose buffering capacity can be systematically quantified through controlled metabolite overfeeding. By mapping metabolite-specific failure thresholds alongside metabolomic, transcriptomic, and proteomic responses, we provided a quantitative framework for understanding how cells tolerate and ultimately fail under conditions of acute metabolite excess.

Failure thresholds across metabolites were highly reproducible, indicating that metabolite toxicity is governed by metabolite-specific buffering capacities rather than a generic stress response. Growing inhibition emerging gradually prior to threshold failure reflects how cellular buffering mechanisms are progressively engaged, indicating that metabolostasis operates through graded regulatory responses rather than binary failure mechanisms. The coincident appearance of amyloid-like assemblies near failure thresholds raises the possibility that metabolite aggregation represents a physical manifestation of metabolostasis breakdown. The precise composition of these assemblies also remains to be determined, as ProteoStat reactivity may reflect co-aggregation of cellular proteins alongside specific metabolite assemblies, as indicated by the reaction with directed antibodies. Direct mechanistic experiments will be required to resolve the extent to which metabolite self-assembly contributes to growth inhibition under overfeeding conditions.

Our analysis further indicates that metabolite tolerance is associated with both metabolic network architecture and physicochemical properties. Metabolites with broader pathway participation exhibited higher failure thresholds and smaller relative pathway disturbances upon overfeeding. These findings suggest that distributed network connectivity provides buffering capacity against metabolite accumulation. In parallel, metabolite solubility was positively associated with intracellular abundance and tolerance, indicating that metabolite chemistry influences the regulatory systems required to maintain homeostasis. Together, these observations suggest that metabolostasis emerges from an interaction between metabolic network topology and metabolite physicochemical properties. More centrally connected metabolites may benefit from distributed buffering across multiple pathways, whereas metabolites with limited connectivity may rely on fewer regulatory routes and therefore exhibit lower tolerance to accumulation.

Integration of matched transcriptomic and proteomic measurements revealed a hierarchical regulatory architecture underlying metabolite resilience. Across metabolite overfeeding conditions, cells consistently engaged a global resource-conservation program characterized by repression of energetically costly anabolic processes. This systemic austerity likely reallocates biosynthetic capacity and energy toward stress defense, providing a general preparedness layer that supports survival under diverse metabolic stresses. Superimposed on this global response were high-magnitude, metabolite-specific regulatory events that activate pathways capable of detoxifying or rerouting excess metabolites. These responses appear tailored to the biochemical properties and metabolic context of individual metabolites, suggesting that cells combine generalized resource management with targeted detoxification strategies. These findings support a two-tiered model of metabolostasis in which a shared preparedness layer establishes baseline resilience, while metabolite-specific adaptive modules buffer acute toxic flux.

An important interpretive consideration is that exogenous amino acid supply in the yeast cells is expected to shift metabolism from de novo biosynthesis toward uptake, reducing NADPH demand and altering the cellular redox balance. The consistent down-attenuation of amino acid biosynthesis genes across overfeeding conditions (**Figure 4E**) may therefore reflect, at least in part, a regulatory adjustment to reduced biosynthetic demand rather than a stress response per se. In parallel, amino acid catabolism can reroute carbon into central metabolism through the Ehrlich pathway, and proteomic signatures related to respiration and oxidative metabolism under overfeeding may reflect this adaptive metabolic rerouting rather than cellular damage. This interpretation is supported by the observation that Ehrlich pathway components, including ARO9 and ARO10, are among the strongest adaptive responses in the T:P analysis, and that both aromatic catabolism and phosphate metabolism genes show consistent regulatory engagement across multiple amino acid overfeeding conditions.

Genetic perturbations provided functional support for this hierarchical buffering architecture. Disruption of metabolite-specific modules impaired tolerance selectively under high metabolite burden, consistent with an adaptive buffering role that becomes engaged once metabolite levels exceed baseline homeostatic capacity. In contrast, perturbation of shared preparedness pathways broadly reduced resilience, indicating that these systems contribute to general stress tolerance across metabolite conditions. In addition, polyphosphate suppressed Phe aggregation kinetics in vitro, suggesting that cellular resilience mechanisms may operate not only through metabolic regulation but also by modulating the physicochemical landscape of metabolite self-assembly. These findings place polyphosphate metabolism at the interface of metabolic regulation and physicochemical stress defense.

Pairwise metabolite overfeeding further revealed synergistic toxicity, hinting that metabolostasis failure thresholds defined for single metabolites may underestimate cellular vulnerability in physiological contexts where multiple metabolites accumulate simultaneously. Potential mechanisms for this synergy include competition for shared buffering pathways, co-aggregation between structurally compatible metabolites, or additive disruption of overlapping metabolic networks. Systematic combinatorial analyses will be necessary to determine the generality and predictability of these interactions and to identify structural or network features that govern metabolite compatibility in co-toxic effects ^42^.

Several limitations of the current study warrant consideration. First, single-metabolite overfeeding in a given yeast strain is a controlled but simplified model that does not fully recapitulate the complexity of physiological metabolite accumulation, where multiple metabolites fluctuate simultaneously and cell-type-specific regulatory architectures may apply. Second, the current analysis is restricted to amino acids. Whether the associations between network topology, chemical solubility, and failure thresholds generalise to other metabolite classes, including organic acids, nucleotides, and lipids, remains to be determined. Third, the aggregation data are based on ProteoStat fluorescence and metabolite-specific antibody imaging, both of which have limitations in specificity for exclusive metabolite versus mixed metabolite-protein aggregates. More direct characterisation of aggregate composition will be important for mechanistic interpretation. Finally, while the two-tiered regulatory architecture is supported by genetic experiments for specific pathway components, comprehensive validation across the full set of predicted buffering nodes remains a priority for future work.

Overall, this work characterizes metabolostasis as a systems-level property emerging from the interplay between metabolic network structure, metabolite physicochemical properties, and post-transcriptional regulatory architecture. The failure thresholds defined here provide quantitative parameters for describing cellular tolerance to metabolite excess, while the two-tier regulatory organization suggests that metabolic resilience is structured through a combination of global preparedness and metabolite-specific adaptive responses. These findings provide a conceptual framework for understanding how cells cope with metabolic perturbations and may have implications for conditions characterized by chronic metabolite accumulation, including inborn errors of metabolism. More broadly, the metabolostasis framework complements existing models of proteostasis and metabolic regulation by highlighting quantitative features of metabolic resilience that may guide future mechanistic and translational studies.

## Methods

### Strains and culture

Yeast strains used in this study and their genotypes are listed in **Table S1**. All strains were confirmed by polymerase chain reaction (PCR). To prevent potential background effects of other metabolites in the synthetic defined media, all growth experiments were performed in minimal media (4 aa) containing only the metabolites required by auxotrophy. The 4aa medium was prepared with 20 g/L glucose and 6.7 g/L Difco yeast nitrogen base without amino acids (Becton, Dickinson and Company, Sparks, MD, USA). The four essential amino acids were supplemented according to the Hartwell formulation^43^, to obtain the following concentrations: Uracil (U0750) 21.4 mg·L^−1^, Histidine (H8125) 21.4 mg·L^−1^, Methionine (M9625) 21.4 mg·L^−1^, and Leucine (L8000) 85.6 mg·L^−1^. The metabolites were supplemented at the concentrations designated for the overfeeding experiments (described below). All chemicals were obtained from Sigma-Aldrich (Rehovot, Israel).

### Kinetic growth assays and failure threshold calculations

Overnight cultures were diluted to OD_600_=0.01 in a final volume of 200 µL per well in a 96-well polystyrene microplate (flat-bottom, sterile; Greiner Bio-One, Kremsmünster, Austria). The plate was incubated at 30 °C with continuous shaking, and OD_600_ measurements were recorded using a Tecan SPARK 10M plate reader (Männedorf, Switzerland). Each assay was performed in triplicates, and the results represent three independent biological replicates.

To determine the failure threshold, the slopes for each growth curve at each of the conditions were first calculated using linear least-square regression with a rolling window of five time points^44,45^. The maximal slope for each condition was then identified and plotted against the concentrations examined. A minimal baseline slope was then defined as 25% of the highest slope. The failure threshold was defined as the average of the two concentrations that fall in the first and second quartiles of the maximal slopes. The calculation was repeated based on at least four biological repeats to determine the mean failure threshold for each metabolite. Herein, the term “failure threshold” refers to the extracellular metabolite concentration at which growth inhibition becomes evident and was operationally defined as the failure threshold of cellular tolerance. It reflects the concentration at which intracellular buffering and regulatory mechanisms become insufficient to maintain growth. Several metabolites did not have significant dose-dependent toxicity, and thus no failure threshold was calculated. For example, as tyrosine is rather insoluble in water^46^, the maximal concentration examined was 2 mM **(Figure S1J)**; leucine and histidine overfeeding was moderately toxic, yet the toxicity pattern was not dose-dependent **(Figure S1D,Q)**; aspartic acid, uracil, cytosine, and adenine also did not cause notable toxicity **(Figure S1S,V-X)**; and the maximal toxicity levels of asparagine and glutamine at their solubility limits (200 mM) were too low to meet the defined criteria of the failure threshold calculation. Thus, failure thresholds were only obtained and presented for 15 metabolites in Figure 1B.

### Metabolite overfeeding

Metabolite concentrations tested were optimized to about 50-60% of the failure threshold on average, to enable sufficient growth of the culture: phenylalanine 8 mM, threonine 8 mM, isoleucine 6 mM, glycine 150 mM, tryptophan 15 mM, valine 15 mM, cysteine 0.5 mM, alanine 13 mM, and methionine 10 mM. The cells were grown in minimal media (4aa), each supplemented with one of the amino acids above at a time. Cellular amino acid composition was analyzed by targeted metabolomics analysis using LC-MS/MS (described below). Specifically, the concentrations of the 20 coding amino acids, as well as ornithine and citrulline, were measured.

### Phe titration in *aro4*Δ yeast strains

To assess the limits of metabolic homeostasis, we reanalyzed Phe titration experiments in *aro4*Δ and WT yeast^32^ by calculating the Area Under the Curve (AUC) of each dose-response growth curve at each Phe concentration (between 0-1.5mM).

### ProteoStat staining and flow cytometry analysis

All ProteoStat staining experiments were performed on live yeast cells. ProteoStat staining was used as a correlative indicator of intracellular amyloid-like aggregation, serving as a general reporter of structured intracellular assemblies. Yeast cultures (5 mL) were grown to mid-log phase (OD_600_≈0.6) and 4 mL was harvested by centrifugation (4,000 rpm, 3 min). Cells were washed in PBS, resuspended in 250 µL PBS, and sonicated for 5 min at 43 kHz. For each condition, 50 µL of the cell suspension was mixed with 450 µL of 1:1000 diluted ProteoStat solution. Cells were incubated in the dark for 15 min at room temperature. Flow cytometry was conducted using the Stratedigm S1000Exi software (San Jose, CA, USA) using a 488 nm laser source and FITC (530/30) and PE-Cy5 (676/29) emission channels, acquiring 50,000 events per sample. Analysis of flow cytometry distributions (**Figure S8**) was performed using FlowJo (Becton-Dickinson).

### Antibody generation

Threonine and isoleucine at a final concentration of 15 mg/mL were dissolved in PBS at 90 °C and then gradually cooled overnight to promote the formation of fibrils as previously reported for isoleucine^14^ and extended here to threonine (**Figure S3**). The fibrillar structures from each metabolite were used as antigens for immunization in rabbits. The immunization was performed five times, followed by polyclonal IgG antibody purification using a protein A column. The antibodies were supplied by Sigma Aldrich (Rehovot, Israel). Phenylalanine antibody was generated as previously described^8^.

### Indirect immunofluorescence staining in yeast

Yeast cultures (4 mL, OD_600_ ≈ 0.6) grown in the presence of 12mM of the indicated metabolites (Phe/ Thr/ Ile) were fixed with 500 µL 37% formaldehyde and 500 µL 1 M KPO₄, followed by 2 h incubation at RT with gentle agitation. Cells were pelleted (4,000 rpm, 3 min), washed 3 times with SP (1.2 M sorbitol, 0.1 M KPO_4_), resuspended in SPβ (SP with 20 mM β-mercaptoethanol) with buffer, and treated with 5 µL Zymolase (12.5 mg/ml in SPβ) at 37 °C for 1.5 h to digest cell walls. Spheroplasts were washed with SP, resuspended in 100 µL SP, and stored at 4 °C overnight.

Poly-L-lysine (200 µL, 0.1%) was applied to an 8-well slide and incubated for ∼60 min at RT. After drying the slide, 100 µL spheroplasts were added to each well and incubated for 60 min. Wells were washed with PBS (200 µL) and blocked with PBT (100 µL) for 2 h at RT. Primary antibody (1:200 in PBT) was added and incubated for 2 h, followed by 6 washes with PBT. Secondary antibody (Cy3-conjugated goat anti-rabbit, 1:200 in PBT) was added and incubated for 1.5 h in the dark, followed by 6 washes with PBS. After disassembling the slide and drying, 10 µL Vectashield mounting medium with DAPI (4′,6-diamidino-2-phenylindole) was applied, covered with a coverslip, and incubated overnight at RT.

Imaging was performed using a Leica TCS SP8 laser confocal microscope with x63/1.4 NA oil immersion objective, using 405 nm and 561 nm excitation lasers and 415-482 nm and 571-648 nm emission photomultipliers for DAPI and Cy3, respectively.

### Metabolite quantification via liquid-chromatography tandem mass spectrometry (LC-MS/MS)

*Metabolite extraction:* Yeast cultures (5 mL) were grown to mid-log phase (OD_600_≈0.6) and 4 mL was quenched with 16 mL ice-cold buffer 1 (60% methanol, 10 mM tricine, pH 7.4, -20 °C). After a 5-min ice incubation, cells were pelleted (1,000 g, 3 min, 4 °C), resuspended in 1 mL cold buffer 1, and pelleted again. The supernatant was removed, and cells were resuspended in 1 mL cold buffer 2 (75% ethanol, 0.5 mM tricine, pH 7.4). Metabolites were extracted by heating at 80 °C for 3 min, followed by cooling on ice for 5 min. After centrifugation (20,000 g, 1 min, 4 °C), 0.9 mL of the supernatant was collected, centrifuged again for 10 min, and 0.8 mL of the cleared extract was stored at -80 °C until analysis.

*LC-MS/MS:* Each sample (10 μL) was mixed with 10 μL of internal standard (Stable Isotope Labeled Amino Acid Mixture, Sigma-Aldrich), agitated for 15 min, derivatized using the AccQTag method, and filtered through 0.2 μm PES nanoFilter vials (Thomson). Quantification was done using a standard curve (0.01-10 μM). The LC-MS/MS system comprised an Acquity I-Class UPLC and a Xevo TQ-S triple quadrupole mass spectrometer (Waters) with chromatographic separation on a UPLC HSS T3 column (100 × 2.1 mm i.d.) and a 1 μL injection volume, as previously described. Data analysis was performed using MassLynx and TargetLynx software (v.4.2, Waters). Analyses were conducted at the Weizmann Institute’s untargeted metabolomics unit. Results represent the average of at least four biological replicates.

*Data analysis:* Replicate-level log₂ fold changes (log₂FC) were calculated by comparing individual metabolite values in treatment samples to the mean abundance of the corresponding metabolite in WT samples. Statistical significance was assessed using two-sample t-tests. Resulting p-values were adjusted for multiple testing using the Benjamini-Hochberg false discovery rate correction to obtain q-values.

### Metabolite regulatory network

We constructed a directed causal metabolite regulatory network using differential metabolite responses and metabolite–metabolite correlations. First, significantly differential metabolite expression (qval < 0.05) upon metabolite overfeeding were filtered for. For each treatment, we defined directed perturbation edges pointing from the treatment to each significantly altered metabolite. Edge weights were computed as ∣log_2_FC∣*(1-q) with signs of log_2_FC encoded as edge-direction signs. To account for metabolite-metabolite dependencies for non-treated metabolites, we next processed replicate-level log_2_FC data. Missing values within each treatment group were median-imputed and metabolite profiles were scaled prior to computing pairwise Pearson correlations. Untreated metabolite–metabolite edges were added for correlations with ∣r∣ > 0.75, retaining the sign of the correlation and removing self-loops. To avoid duplicating direct perturbation edges, correlation edges originating from treated metabolites that were excluded. Perturbation edges and correlation edges were then normalized and combined. Direct perturbation edges were assigned 60% of total edge weight (α = 0.6), with the remaining weight assigned to correlation edges to capture endogenous metabolic structure. The network was constructed in *igraph* and visualised with edge color indicating sign (red = positive, blue = negative) and line width proportional to normalized weight. Network influence of metabolites in the network were calculated using personalized-page rank.

### Metabolite properties

Aqueous solubility of metabolites was obtained from PubChem. Hydrophobicity and ß-sheet propensities were obtained from CamSol^47^. Aggregation propensities at pH2, pH7, and pH13 were obtained from published literature^34^. All collated metabolite properties used in this study are available in **Table S2**.

### Next-generation sequencing (NGS)

#### RNA extraction and quality control

Yeast cultures (5 mL) were grown to mid-log phase (OD_600_≈0.6), and 1 mL of cells was harvested by centrifugation (10,000 g, 1 min) and washed twice with ice-cold PBS. Pellets were snap-frozen in liquid nitrogen and stored at -80 °C. RNA was extracted using the MasterPure Yeast RNA purification kit, followed by DNase I treatment for 30 min. RNA aliquots (5-10 µL) were stored at -80 °C. Concentration was determined using the Qubit RNA High-Sensitivity assay kit (Invitrogen). The RNA quality was assessed using the Agilent 4200 TapeStation, ensuring RNA Integrity Numbers (RIN)>8.

#### Library preparation and NGS

cDNA libraries were generated from 300 ng of total RNA using the NEBNext Ultra II Directional RNA Library Prep Kit, with the NEBNext Poly(A) mRNA Magnetic Isolation Module (NEB #E7760), according to manufacturer’s instructions. Amplification was performed by 15 PCR cycles with the NEBNext Multiplex Oligos kit (NEB #E6440). The quality and quantity of each library were assessed using the TapeStation 4200 system (Agilent) and a Qubit 3.0 Fluorometer (Invitrogen). Samples were pooled and sequenced on the NextSeq 1000/2000 P2 Kit (100 Cycles) (Illumina #20046811) using a 100 bp single-read output run on a NextSeq 2000 system (Illumina). Library preparation and sequencing were performed as a service by the Genomics Research Unit at the Life Sciences Cancer Research Core Facility, Tel-Aviv University.

#### Data processing

The FASTQ files were analyzed using Partek Flow build version 10.0.21.0411. Reads were trimmed to remove low-quality bases from the 3’ end (Phred score<20). The RNA-seq reads were mapped to the *S. cerevisiae* reference genome (GCF_000146045.2, R64) obtained from the NCBI Genome Assembly database. Alignment was performed using STAR^48^ (v2.7.8a). Reads were quantified using Partek E/M^49^. A total of 6,006 genes were detected and statistically analyzed using *DESeq2* median ratio normalization^50^. Differentially expressed genes were identified with a false discovery rate (FDR) threshold of <0.05, between each treatment and the control. The raw data analysis was performed as a service at the Bioinformatics Unit, Faculty of Life Sciences, Tel Aviv University.

### Global proteome analysis

#### Protein extraction

Yeast cultures (5 mL) were grown to mid-log phase (OD_600_≈0.6), and 1 mL of cells was harvested by centrifugation, washed twice with ice-cold PBS, snap-frozen in liquid nitrogen, and stored at -80 °C. For extraction, pellets were lysed with 200 µL protein extraction buffer (50 mM Tris, 5% SDS, pH 7.6) and glass beads by bead beating for 8 cycles (1 min at 4 °C, with 1-min intervals on ice). Lysates were boiled (90 °C, 5 min), placed on ice (5 min), separated from beads using a pipette, and stored at -80 °C until analysis.

#### Sample preparation

The samples were prepared for mass spectrometry analysis as previously described. Briefly, lysates were incubated at 96 °C for 5 min, followed by six cycles of 30 seconds of sonication (Bioruptor Pico, Diagenode, USA). Protein concentration was measured using the bicinchoninic acid (BCA) assay (Thermo Fisher, USA), and a total of 25 μg of protein was reduced with 5 mM dithiothreitol and alkylated with 10 mM iodoacetamide in the dark. Phosphoric acid was added to the lysates to a final concentration of 1.2%, followed by a 90:10% mixture of methanol/50 mM ammonium bicarbonate. Each sample was loaded onto an S-Trap microplate (Protifi, USA) and digested with trypsin at a 1:50 trypsin/protein ratio for 1.5 h at 47 °C. The digested peptides were eluted with 50 mM ammonium bicarbonate, with additional trypsin added to this fraction, and incubated overnight at 37 °C. Two additional elution steps were performed using 0.2% formic acid and 0.2% formic acid in 50% acetonitrile. The three elution fractions were pooled together and vacuum-centrifuged to dryness. Samples were stored at -80°C until analysis.

#### Liquid chromatography

ULC/MS grade solvents were used for all chromatographic steps. Each sample was loaded using split-less nano-Ultra Performance Liquid Chromatography (NanoElute2, Bruker Daltonics). The mobile phases were: (A) water with 0.1% formic acid and (B) acetonitrile with 0.1% formic acid. Peptides were separated using an Aurora column (75 µm ID x 25 cm, IonOpticks) at a flow rate of 0.3 µL/min. The elution gradient was as follows: 2% to 30% B over 41 min, 30% to 90% B over 2 min, maintained at 90% B for 3 min, and then returned to the initial conditions.

#### Mass spectrometry

The nanoUPLC was coupled online through a 10 µm ZDV emitter to a timsTOF PRO mass spectrometer (Bruker Daltonics). Data were acquired in parallel accumulation–serial fragmentation combined with data-independent acquisition (DIA-PASEF) mode. For ion mobility, the 1/K0 range was set to 0.67-1.41 V/cm², with a ramp time of 100 milliseconds and an estimated cycle time of 1.48 seconds. A total of 36 windows, each 25 Da wide with a 1 Da overlap, were set over a mass range of 319.5-1184.5 Da.

#### Data processing

Raw data were processed using Spectronaut v18.1 (Biognosys). The data were searched against the SwissProt *S. cerevisiae* proteome database (January 2024 version, 6,060 entries) and included a common contaminants list. Carbamidomethylation of cysteine (C) was set as a fixed modification. Oxidation of methionine (M), deamidation of asparagine (N) and glutamine (Q), and protein N-terminal acetylation were set as variable modifications. Protein intensities were used for further calculations in Perseus v1.6.2.3. Decoy hits were filtered out. Protein intensities were log_2_-transformed, and only proteins with at least two valid values in at least one experimental group were retained. The remaining missing values were imputed using a random low-range distribution. Student’s t-tests were performed between the relevant groups to identify significant changes in protein levels. To ensure the accuracy of the results, all subsequent analysis steps were performed only on proteins identified by at least two peptides. The global proteome analysis was performed as a service at The De Botton Protein Profiling Institute of the Nancy and Stephen Grand Israel National Center for Personalized Medicine, Weizmann Institute of Science.

### Post-transcriptional (T:P) regulatory analysis

Matched transcriptomic and proteomic profiles (4408 genes/proteins) from the same cultures across all metabolite treatments were used for the T:P regulatory analysis. To assess post-transcriptional regulation under metabolite overfeeding, genes were classified based on the concordance between mRNA and protein fold changes (log2FC). All significantly altered genes (*Genes_ALL_*) at either the transcriptomic or proteomic level were included in the total gene set. Where *Attenuation_D_* = number of down-attenuated genes (log_2_FC(mRNA) < 0 & log_2_FC(protein)-log_2_FC(mRNA) > 0.3); *Attenuation_U_* = number of up-attenuated genes (log_2_FC(mRNA) > 0 & log_2_FC(protein)-log_2_FC(mRNA) < -0.3); *Gain_U_* = number of up-gain genes (log_2_FC(mRNA) > 0 & log_2_FC(protein)-log_2_FC(mRNA) > 0.3); *Gain_D_* = number of down-gain genes (log_2_FC(mRNA) < 0 & log_2_FC(protein)-log_2_FC(mRNA) < 0.3); and *Gain_Ntrl_* |log_2_FC(protein)-log_2_FC(mRNA)| < 0.3). A threshold of |Δ| = 0.3 was implemented to account for noise in fold-change differences between transcript and protein levels.

To quantify cellular adaptive strategies, we introduce the following metrics:

1. **Adaptive Strategy Preference (ASP)** quantifying the ratio of resource-conservation to amplification strategies.

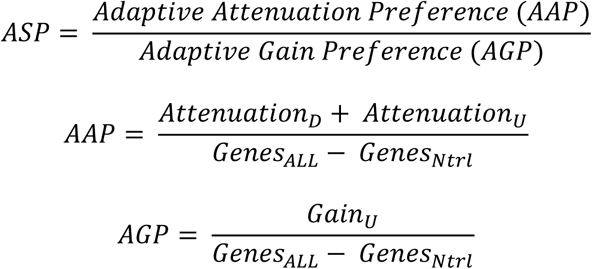

1. **Directional Bias** defined as the ratio of repression-oriented events to induction-oriented events.

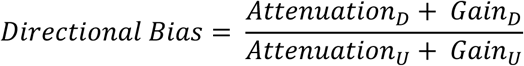

1. **Total T:P Control Intensity (TTCI)** quantifying the total post-transcriptional regulatory magnitude.

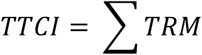

where,

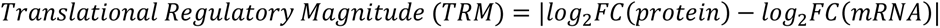

All analyses were performed on n = 9 metabolite overfeeding conditions and are available in the **Supplementary Data**. Correlations between ASP, AAP, Directional Bias, and TTCI with metabolite failure thresholds were assessed using Pearson correlation.

### Specificity of Up-attenuated genes

To quantify the specificity of ‘buffering’ genes to metabolite perturbation, we computed the number of perturbation conditions each gene was significantly affected in. For each metabolite condition, up-attenuated genes were ranked by TRM score. The top 200 genes for each perturbation condition were used for this calculation. To compare the average specificity of specific functional groups, we took the average of the number of conditions a family of genes (eg. Aro(s)/Pho(s)/Vtc(s)) were found to be significantly affected by.

### Quantification of yeast growth and fractional inhibition (ARO, PHO, VTC mutants & metabolite overfeeding experiments)

Yeast growth was quantified by calculating the AUC of the dose-response yeast growth curves (up to 35h). Fractional inhibition for each treatment was calculated relative to the relevant control:

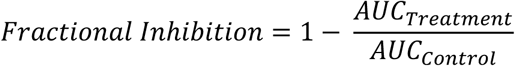

### PolyP in vitro

PolyP was kindly provided by Dr. Toshikazu Shiba (Regenetiss, Japan). Phe aggregation assays were performed using 35 mg ml⁻¹ Phe dissolved in phosphate-buffered saline (PBS). To assess the effect of PolyP, short-chain polyP was added to Phe solutions at final concentrations of 15, 45, or 60 mM following a 3-hour pre-incubation period allowing for partial aggregation. Reaction mixtures (200 µl) were transferred to 96-well plates and incubated for 20 h at 30 °C. Turbidity measurements monitored aggregation at 405 nm. For quantification, turbidity values at 20 h were normalized to the Phe-only control.

### Combinatory effects of metabolite toxicity

Phe, Ile, and Thr were dissolved in SD 4aa medium to generate single-amino-acid solutions at 4 mM and 8 mM. Amino acid combinations were prepared by dissolving each amino acid at 4 mM in the same medium.

To evaluate whether combinations of metabolites exhibited synergistic effects beyond what would be expected under non-interacting conditions, we applied the Bliss Independence^41^ model as a null reference framework. Under this probabilistic model, the expected fractional inhibition of a combination of two treatments is calculated from the individual effects of each treatment acting alone, assuming they act through independent mechanisms. Observed combination effects that exceeded the Bliss-predicted values were interpreted as synergistic interactions.

## Acknowledgments

This project was made possible through the support of a grant from the John Templeton Foundation (grant number 61735) and Israel Science Foundation (grant number 969/24). The opinions expressed in this publication are those of the authors and do not necessarily reflect the views of the John Templeton Foundation. We thank the members of the Gazit laboratory for helpful discussions. We would like to thank Maya Schuldiner for providing strains and for her valuable insights and to Meital Kupervaser and The De Botton Protein Profiling Institute of the Nancy and Stephen Grand Israel National Center for Personalized Medicine, Weizmann Institute of Science, for assistance in protein profiling. We thank Tamar Katzir, Karin Smorodinsky-Atias, and Hila Kobo from the Genomics Research Unit at the Life Sciences Cancer Research Core Facility, Tel Aviv University, for assistance in mRNA sequencing. Special thanks to Alexandra Lichtenstein for assistance with confocal microscopy, Orit Sagi-Assif for assistance with flow cytometry, Tevie Mehlman for assistance in LC-MS measurements and George Levi for assistance with TEM imaging. PolyP was kindly provided by Dr. Toshikazu Shiba (Regenetiss, Japan).

## Author contribution

SAL, CML, VM, EG, and DLB-Y conceived the project and designed the experiments. SAL performed most of the experiments with assistance from EAM, HA, MG, KK, ML, IS-B and YD. IS-B and HA assisted in antibody generation. ML performed the failure thresholds calculations. AB performed the LC-MS measurements. DLB-Y coordinated the work performed by various team members. CML analyzed most of the data with assistance from SAL and MP-C. SAL, CML, MV, EG, DLB-Y wrote the manuscript. VM, EG, and DLB-Y edited the manuscript based on recent advancements in the field.

