## Supplementary Figures for "Metabolostasis failure thresholds are linked with network topology, metabolite solubility, and translational control"

*[1]Shmunis School of Biomedicine and Cancer Research,*
*George S. Wise Faculty of Life Sciences, Tel Aviv University, Tel Aviv 6997801, Israel*
*[2]Centre for Misfolding Diseases, Yusuf Hamied Department of Chemistry,*
*University of Cambridge, Cambridge CB2 1EW, UK*
*[3]Bioinformatics Unit, George S. Wise Faculty of Life Science,*
*Tel Aviv University, Tel Aviv 6997801, Israel*
*[4]Department of Life Sciences Core Facilities, Weizmann Institute of Science, Rehovot, Israel*
[5]Sagol *School of Neuroscience, Tel Aviv University, Tel Aviv 6997801, Israel*

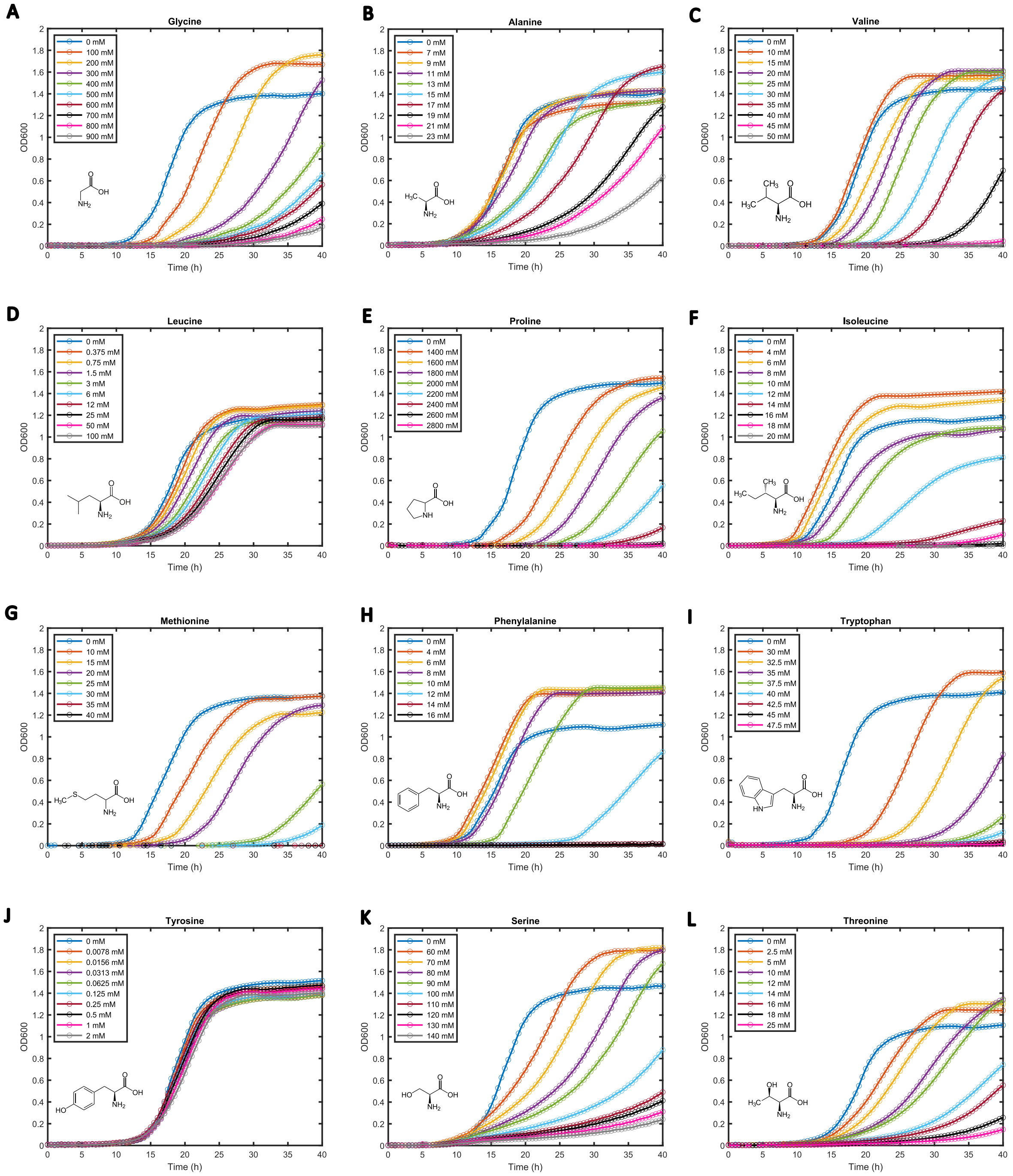
A
Glycine
0 mM
100 mM
200 mM
300 mM
400 mM
500 mM
600 mM
700 mM
800 mM
900 mM
B
Alanine
0 mM
7 mM
9 mM
11 mM
13 mM
15 mM
17 mM
19 mM
21 mM
23 mM
C
Valine
0 mM
10 mM
15 mM
20 mM
25 mM
30 mM
35 mM
40 mM
45 mM
50 mM
D
Leucine
0 mM
0.375 mM
0.75 mM
1.5 mM
3 mM
6 mM
12 mM
25 mM
50 mM
100 mM
E
Proline
0 mM
1400 mM
1600 mM
1800 mM
2000 mM
2200 mM
2400 mM
2600 mM
2800 mM
F
Isoleucine
0 mM
4 mM
6 mM
8 mM
10 mM
12 mM
14 mM
16 mM
18 mM
20 mM
G
Methionine
0 mM
10 mM
15 mM
20 mM
25 mM
30 mM
35 mM
40 mM
H
Phenylalanine
0 mM
4 mM
6 mM
8 mM
10 mM
12 mM
14 mM
16 mM
I
Tryptophan
0 mM
30 mM
32.5 mM
35 mM
37.5 mM
40 mM
42.5 mM
45 mM
47.5 mM
J
Tyrosine
0 mM
0.0078 mM
0.0156 mM
0.0313 mM
0.0625 mM
0.125 mM
0.25 mM
0.5 mM
1 mM
2 mM
K
Serine
0 mM
60 mM
70 mM
80 mM
90 mM
100 mM
110 mM
120 mM
130 mM
140 mM
L
Threonine
0 mM
2.5 mM
5 mM
10 mM
12 mM
14 mM
16 mM
18 mM
25 mM
OD600
Time (h)

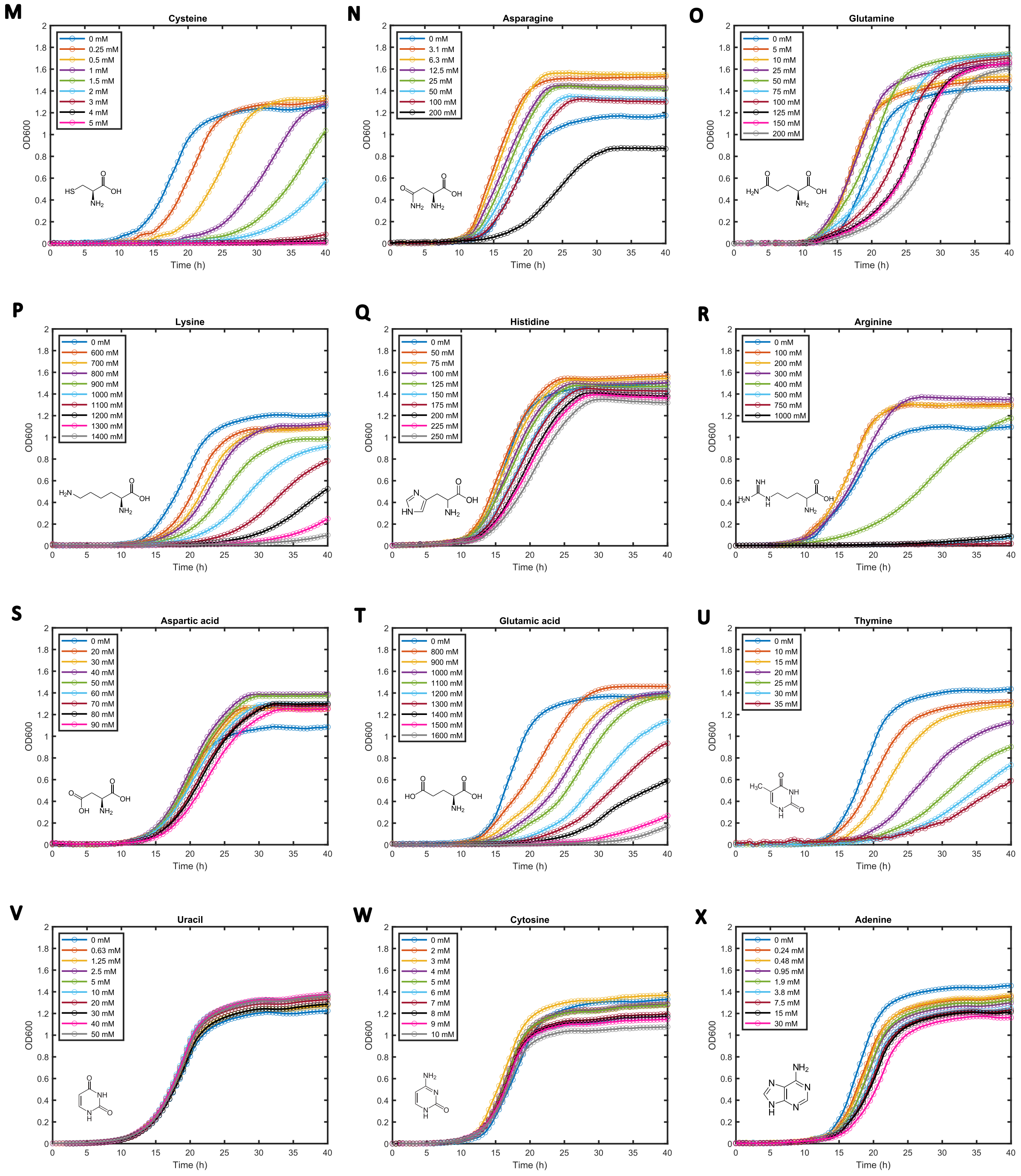


**Figure S1. Dose-response growth curves for individual metabolite overfeeding conditions.** Growth curves of *S. cerevisiae* under increasing concentrations of each metabolite. Results are representative of at least four biological replicates used to calculate failure thresholds (**Figure 1B**). **(A-X)** Metabolites are grouped by chemical properties: nonpolar **(A-G)**, aromatic **(H-J)**, polar (**K-O)**, positively charged **(P-R)**, negatively charged **(S-T)**, and nucleobases **(U-X)**. Twenty-four metabolites were tested; guanine was excluded due to insolubility in water.

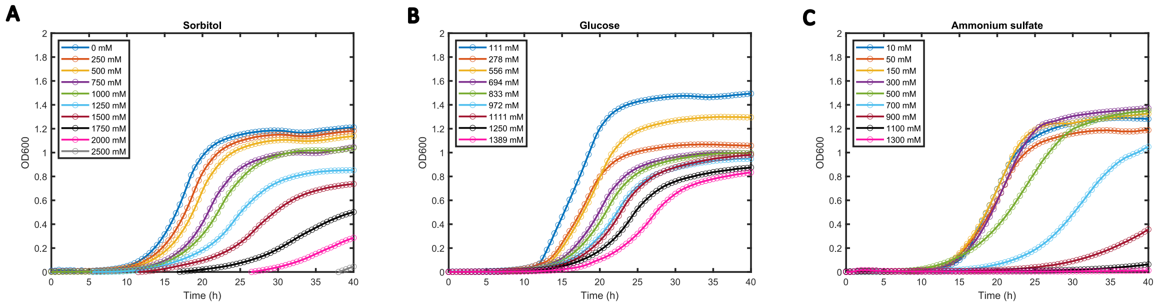


**Figure S2. Non-metabolite-specific stress controls**. (**A-C**) Growth curves of *S. cerevisiae* under increasing concentrations of **(A)** sorbitol (osmotic stress), **(B)** glucose (carbon source excess), and **(C)** ammonium sulfate (nitrogen excess). Significant non-specific growth effects were observed only above 400 mM, confirming that failure thresholds below this concentration reflect metabolite-specific toxicity rather than general osmotic or nutrient stress.

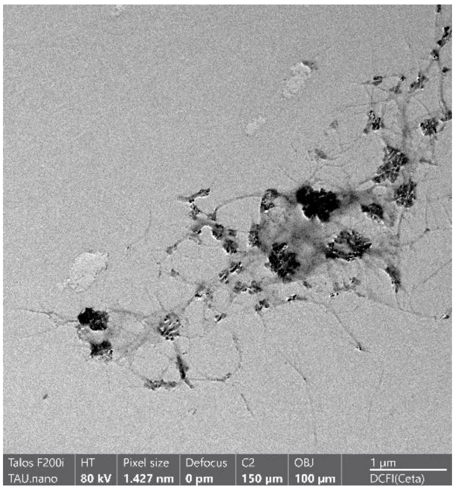


**Figure S3.** Threonine assembles into fibrils. Representative TEM image of Threonine fibrils at 15 mg/mL.

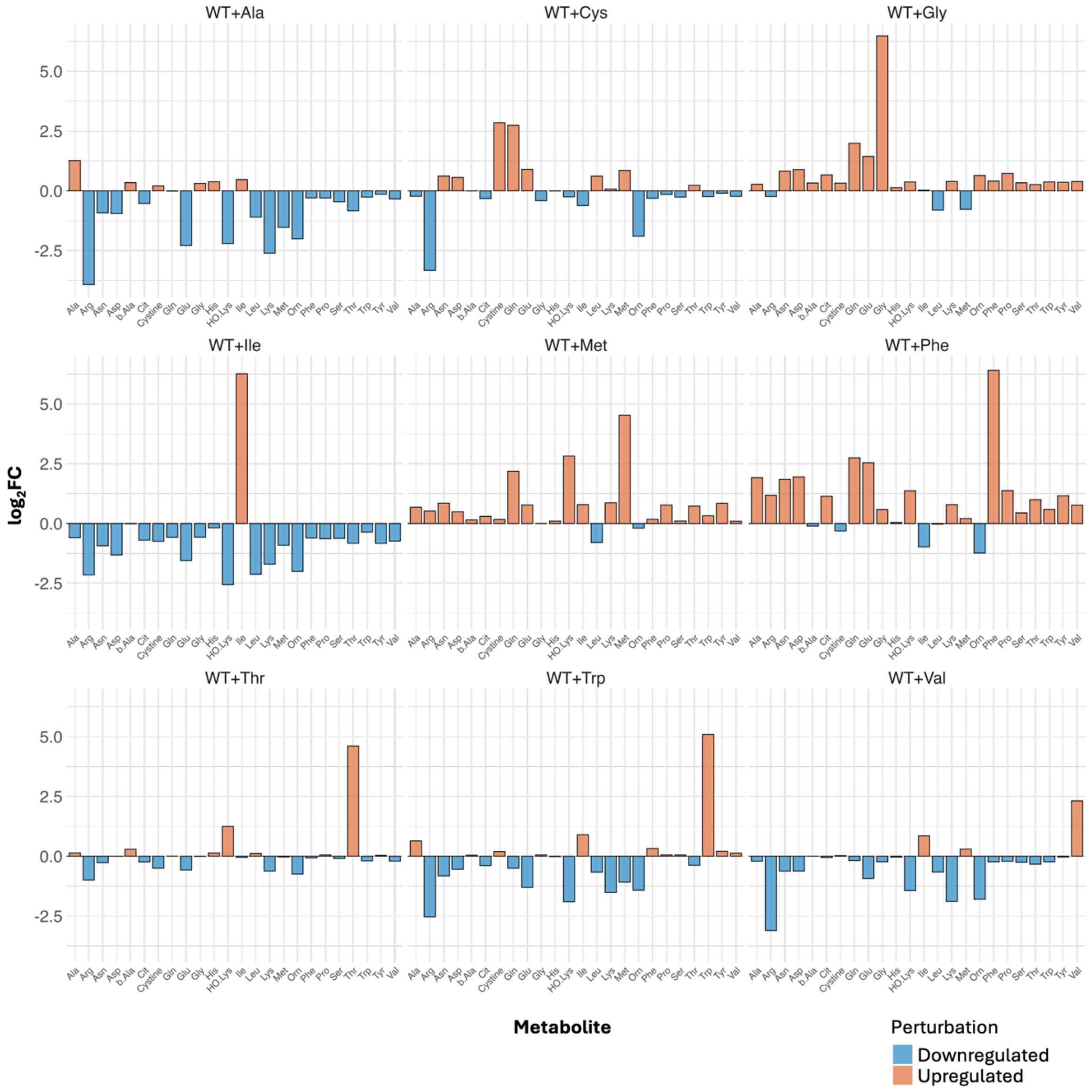


**Figure S4. Intracellular metabolite signatures under overfeeding conditions.** Targeted LC–MS/MS profiling confirms significant cellular uptake of each overfed metabolite. Each treatment also elicited a unique secondary metabolic signature.

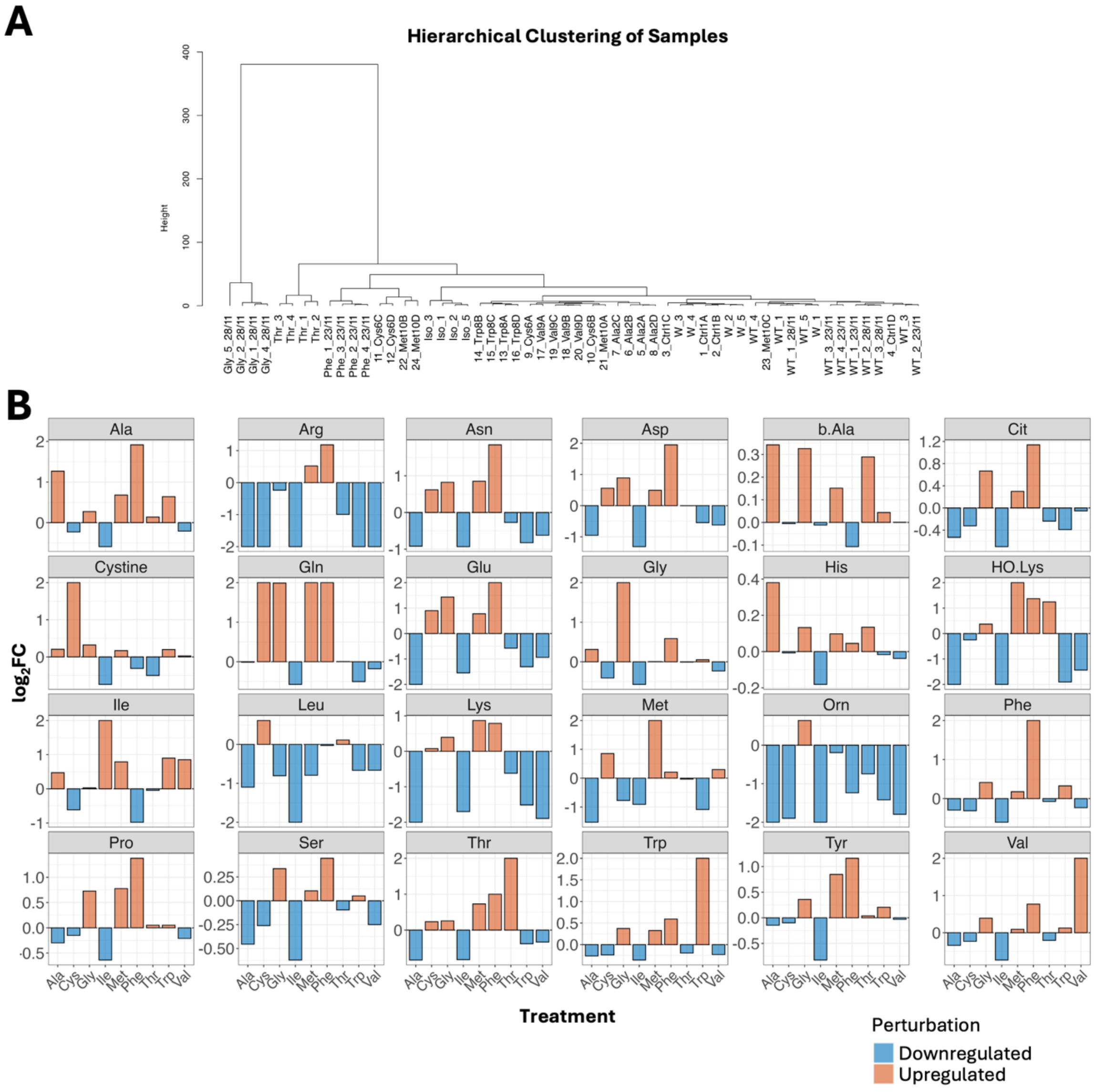


**Figure S5. Condition-specific metabolic shifts. (A)** Evaluation of global metabolome structure across experimental replicates reveals high reproducibility between replicates. Direction and magnitude of metabolic shifts were highly condition-specific. **(B)** Relative abundance of each metabolite under different overfeeding conditions. Targeted analysis of metabolic trends reveals the recurrent depletion of arginine and ornithine across multiple perturbations. This suggests that the urea cycle may act as a primary buffer for acute metabolic imbalance.

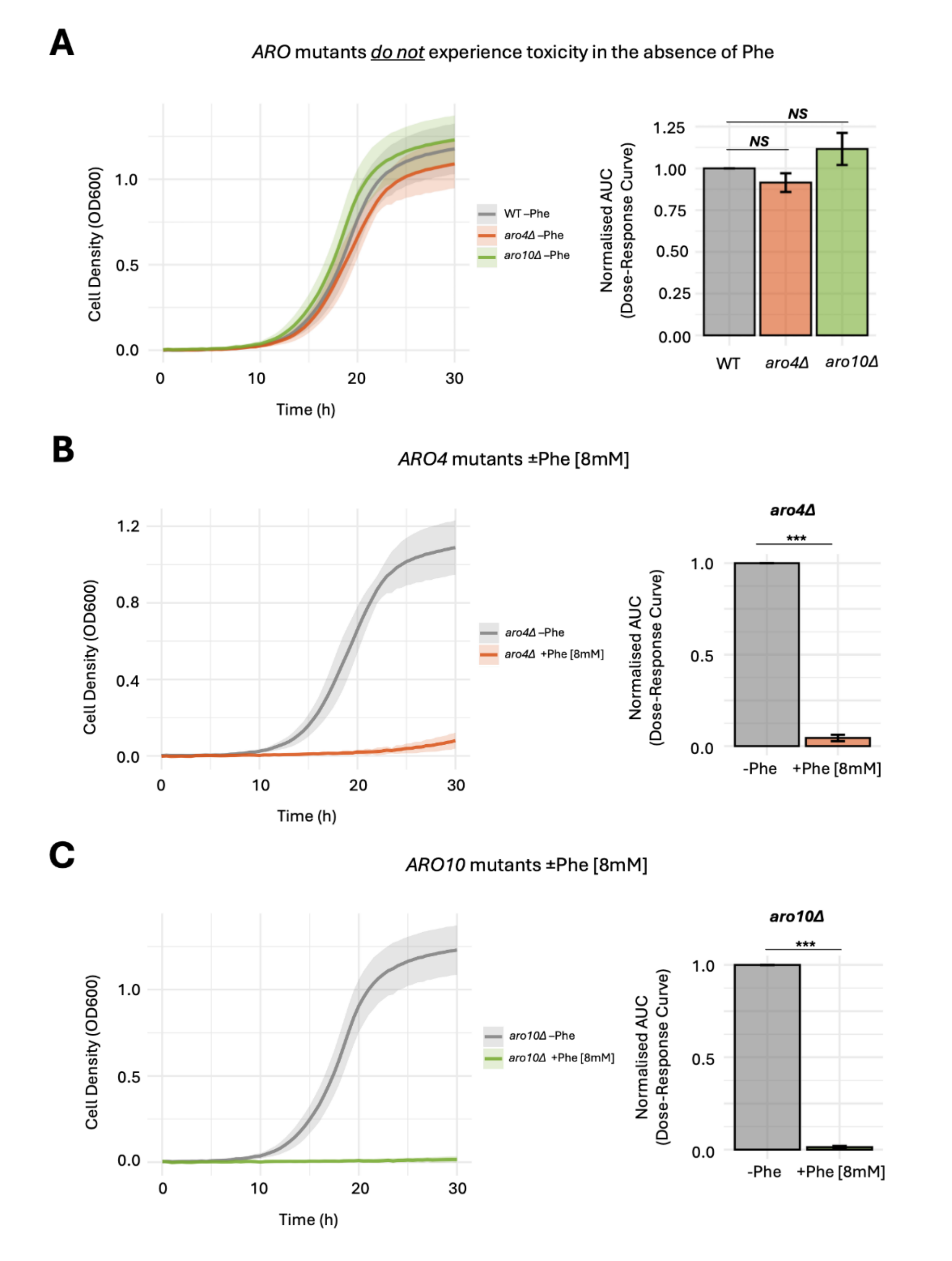


**Figure S6. Control experiments of *ARO* mutants ±Phe. (A)** *aro4Δ* and *aro10Δ* strains do not display toxicity under normal conditions, but only in the presence of Phe. **(B)** *aro4Δ* cells experience toxicity upon treatment with Phe [8mM]. **(C)** *aro10Δ* cells experience toxicity upon treatment with Phe [8mM]. (***) $p \leq 0.001$ and *NS*, not significant.

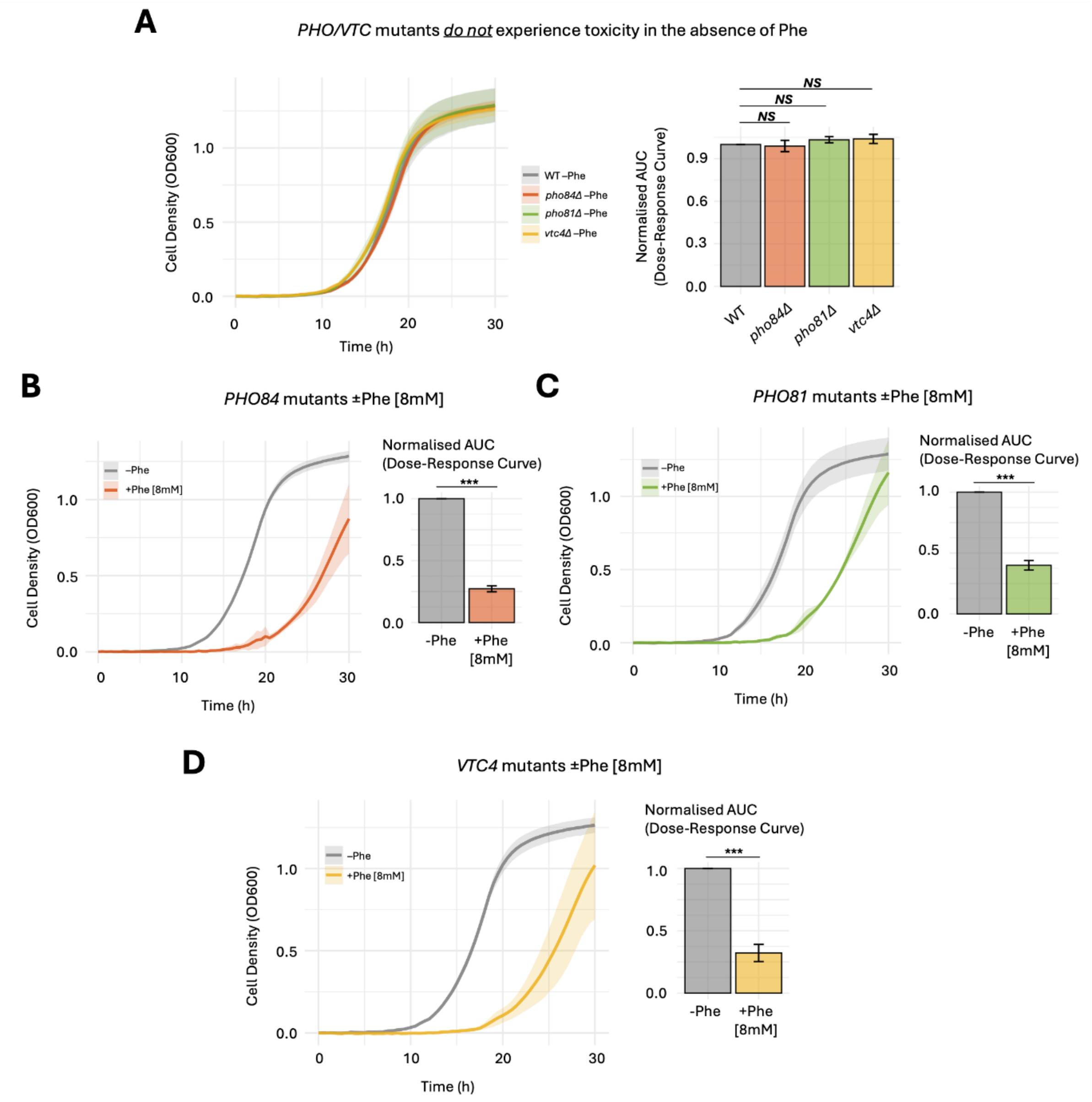


**Figure S7. Control experiments of *PHO/VTC* mutants ±Phe. (A)** *pho84Δ, pho81Δ* and *vtc4Δ* strains do not display toxicity under normal conditions, but only in the presence of Phe. **(B)** *pho84Δ* cells experience toxicity upon treatment with Phe [8mM]. **(C)** *pho81Δ* cells experience toxicity upon treatment with Phe [8mM]. **(D)** *vtc4Δ* cells experience toxicity upon treatment with Phe [8mM]. (***) $p \leq 0.001$ and *NS*, not significant

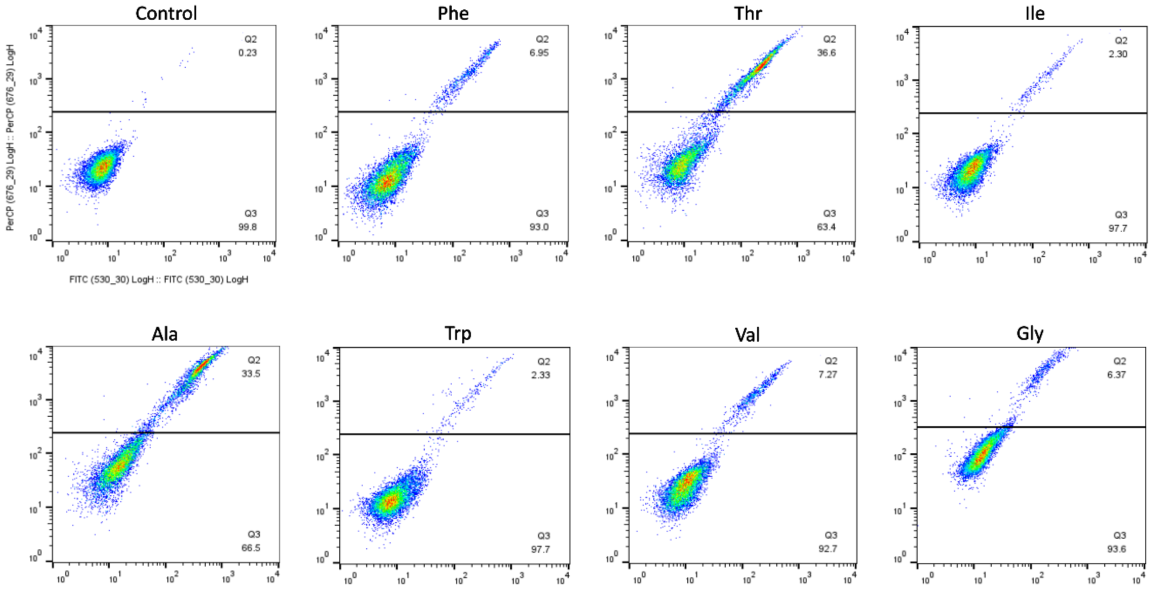


**Figure S8. Representative flow cytometry distributions.** PerCP and FITC values are presented on a log scale.

**Table S1: *Saccharomyces cerevisiae* strains used in this study**. All strains are isogenic derivatives of BY4741. Deletion strains carry the KanMX cassette replacing the indicated open reading frame.

| Yeast strains | Genetic background | Source |
|---|---|---|
| Wild-type, BY4741 | MATa, his3-Δ1, leu2Δ-0, met15-Δ0, ura3-Δ0 | Laor et al.[1], 2019 |
| *aro4Δ* | MATa, his3-Δ1, leu2Δ-0, met15-Δ0, ura3-Δ0, aro4::KanMX | Adsi et al.[2], 2026 |
| *aro10Δ* | MATa, his3-Δ1, leu2Δ-0, met15-Δ0, ura3-Δ0, aro10::KanMX | Deletion library, Giaever et al.[3], 2002 |
| *pho81Δ* | MATa, his3-Δ1, leu2Δ-0, met15-Δ0, ura3-Δ0, pho81::KanMX | Deletion library, Giaever et al.[3], 2002 |
| *pho84Δ* | MATa, his3-Δ1, leu2Δ-0, met15-Δ0, ura3-Δ0, pho84::KanMX | Deletion library, Giaever et al.[3], 2002 |
| *vtc4Δ* | MATa, his3-Δ1, leu2Δ-0, met15-Δ0, ura3-Δ0, vtc4::KanMX | Deletion library, Giaever et al.[3], 2002 |

**Table S2: Physicochemical properties of amino acids used in this study.** Aqueous solubility values were obtained from PubChem. Hydrophobicity and β-sheet propensity scores were obtained from CamSol. Chemical group classifications follow standard amino acid side-chain chemistry.

| Name | Chemical group | Solubility in water | Hydrop hobicity | β-sheet propensity | Aggregation propensity (pH 2) | Aggregation propensity (pH 7) | Aggregation propensity (pH 13) |
|---|---|---|---|---|---|---|---|
| Alanine (Ala) | non-polar | 164 g/L | 41 | 0.97 | -3.31 | -3.31 | -3.31 |
| Arginine (Arg) | positively-charged | 182 g/L | -14 | 0.9 | -11.93 | -11.93 | -11.85 |
| Asparagine (Asn) | polar | 29.4 g/L | -28 | 0.65 | -6.02 | -6.02 | -6.02 |
| Aspartic acid (Asp) | negatively-charged | 5.36 g/L | -55 | 0.8 | -9.42 | -4.38 | -9.42 |
| Cysteine (Cys) | polar | 280 g/L | 49 | 1.3 | 1.61 | 1.61 | -3.44 |
| Glutamic acid (Glu) | negatively-charged | 8.57 g/L | -10 | 0.26 | -10.38 | -6.73 | -10.38 |
| Glutamine (Gln) | polar | 41.3 g/L | -31 | 1.23 | -6 | -6 | -6 |
| Glycine (Gly) | non-polar | 249 g/L | 0 | 0.81 | -3.96 | -3.96 | -3.96 |
| Histidine (His) | positively-charged | 45.6 g/L | 8 | 0.71 | -4.31 | -9.36 | -4.31 |
| Isoleucine (Ile) | non-polar | 34.4 g/L | 99 | 1.6 | 0.93 | 0.93 | 0.93 |
| Leucine (Leu) | non-polar | 21.5 g/L | 97 | 1.22 | -0.25 | -0.25 | -0.25 |
| Lysine (Lys) | positively-charged | 584 g/L | -23 | 0.74 | -9.55 | -9.55 | -9.47 |
| Methionine (Met) | non-polar | 56.6 g/L | 74 | 1.67 | -1.06 | -1.06 | -1.06 |
| Phenylalanine (Phe) | aromatic | 26.4 g/L | 100 | 1.28 | 2.8 | 2.8 | 2.8 |
| Proline (Pro) | non-polar | 162 g/L | -46 | 0.62 | -11.96 | -11.96 | -11.96 |
| Serine (Ser) | polar | 425 g/L | -5 | 0.72 | -5.08 | -5.08 | -5.08 |
| Threonine (Thr) | polar | 97.0 g/L | 13 | 1.2 | -2.12 | -2.12 | -2.12 |
| Tryptophan (Trp) | aromatic | 13.4 g/L | 97 | 1.19 | 2.92 | 2.92 | 2.92 |
| Tyrosine (Tyr) | aromatic | 0.453 g/L | 63 | 1.29 | 1.03 | 1.03 | 1.03 |
| Valine (Val) | non-polar | 58.5 g/L | 76 | 1.65 | 0.49 | 0.49 | 0.49 |

**Supplementary Methods**

**Transmission Electron Microscopy**

Threonine was dissolved at 15 mg/mL at 90 °C in PBS buffer to obtain a homogeneous monomeric solution. The solution was gradually cooled down and incubated overnight at room temperature to facilitate self-assembly. For the TEM images samples were vortexed, and 10 µL of each was drop-cast onto 400-mesh copper grids (Electron Microscopy Sciences, Hatfield, PA, USA). The grids were incubated at room temperature for 5 min, and excess fluid was absorbed. Samples were visualized using a Talos F200i (S)TEM equipped with a field emission gun and a Ceta-M Camera. The accelerating voltage was 80 kV, and the beam current was ~0.5 nA.
